# Exploring the interaction network of a synthetic gut bacterial community

**DOI:** 10.1101/2021.02.25.432904

**Authors:** Anna S. Weiss, Anna G. Burrichter, Abilash Chakravarthy Durai Raj, Alexandra von Strempel, Chen Meng, Karin Kleigrewe, Philipp C. Münch, Luis Rössler, Claudia Huber, Wolfgang Eisenreich, Lara M. Jochum, Stephanie Göing, Kirsten Jung, Alvaro Sanchez, Bärbel Stecher

## Abstract

A key challenge in microbiome research is to predict functionality from microbial community composition. As central microbiota functions are determined by bacterial community networks it is important to gain insight into the principles that govern bacteria-bacteria interactions. Here, we focused on growth and metabolic interactions of the Oligo-Mouse-Microbiota (OMM^12^) synthetic bacterial community, which is increasingly used as model system in gut microbiome research. Using a bottom-up approach, we uncovered the directionality of strain-strain interactions in mono- and pairwise co-culture experiments, as well as in community batch culture. Metabolomics analysis of spent culture supernatant of individual strains in combination with genome-informed pathway reconstruction provided insights into the metabolic potential of the individual community members. Thereby, we could show that the OMM^12^ interaction network is shaped by both, exploitative and interference competition *in vitro.* In particular, *Enterococcus faecalis* KB1 was identified as important driver of community composition by affecting the abundance of several other consortium members. Together, this study gives fundamental insight into key drivers and mechanistic basis of the OMM^12^ interaction network, which serves as knowledge base for future mechanistic studies.

## Introduction

The mammalian gastrointestinal tract harbors hundreds of bacterial species that occupy distinct ecological niches (1, 2). Diversity and stable coexistence of community members after initial assembly result in exclusion of invaders (3, 4). Community assembly and stability are inherently driven by commensal or cooperative trophic interactions, in which metabolic by- or end products of one species are the resource for another one (5–7). At the same time, bacteria compete for substrates by employing diverse predatory mechanisms, like the production of bacteriocins (8). These interaction patterns form complex ecological networks and determine community-level functions of the microbiota including dietary breakdown, metabolite production and colonization resistance (9–11). Consequently, disruption of bacterial networks by antibiotics, disease or diet-mediated interventions results in impairment of community-level functions (12, 13). To be able to predict, preserve and manipulate microbial community function, it is important to identify functionally important members and understand relevant interaction mechanisms between individual bacteria.

A multitude of different approaches have been used to characterize ecological networks of microbial communities. Function-related patterns in native microbial communities can been identified by systems biology approaches, combining metagenomics, metatranscriptomics and metabolomics analyses (14). Together with stable-isotope probing methodologies microorganisms with specific metabolic properties can be identified (15). Potentially interacting species may be predicted from co-occurrence analysis supported by genome guided metabolic modeling (16–18). To experimentally verify the key ecological, structural and functional role of certain species in community structure and function, synthetic microbial consortia provide several advantages over native communities. As they are well-characterized, scalable and experimentally tractable, these systems are increasingly used to gain a mechanistic understanding of gut microbial ecology (19–22).

The Oligo-Mouse-Microbiota (OMM^12^) is a synthetic bacterial community, which stably colonizes mice and provides colonization resistance against enteropathogen infection (23–26). The OMM^12^ comprises twelve bacterial species *(Enterococcus faecalis* KB1, *Limosilactobacillus reuteri* I49, *Bifidobacterium animalis* YL2, *Clostridium innocuum* I46, *Blautia coccoides* YL58, *Enterocloster clostridioformis* YL32, *Flavonifractor plautii* YL31, *Acutalibacter muris* KB18, *Bacteroides caecimuris* I48, *Muribaculum intestinale* YL27, *Akkermansia muciniphila* YL44 and *Turicimonas muris* YL45), representing the five major eubacterial phyla in the murine gastrointestinal tract (27) (**Fig. 1A**). The model is freely available for non-commercial use (28), and is therefore increasingly employed in preclinical microbiome research (29–32). So far little is known about the system’s ecology and metabolic capabilities, both of which are factors that determine assembly, population dynamics and bacterial community functionality. Therefore, we aimed for a comprehensive exploration of the metabolic potential (i.e., substrates, metabolism and end products) and interactions between individual members of the OMM^12^ consortium. We employed a bottom-up approach connecting outcomes of mono- and pairwise co-culture experiments with observations from complex communities in *in vitro* batch culture. Furthermore, we combined metabolomics analysis of spent culture supernatants with genome-informed pathway reconstruction and generated draft metabolic models of the OMM^12^ consortium. Overall, we find that the majority of *in vitro* strain-strain interactions is amensalistic or competitive. In accordance, bacteriocin production and substrate overlap between the individual strains was correlated with negative strain-strain interaction *in vitro*, revealing potentially underlying mechanisms. Together, this work identified key interaction patterns among OMM^12^ strains relevant in community assembly and functionality.

**Figure 1:**
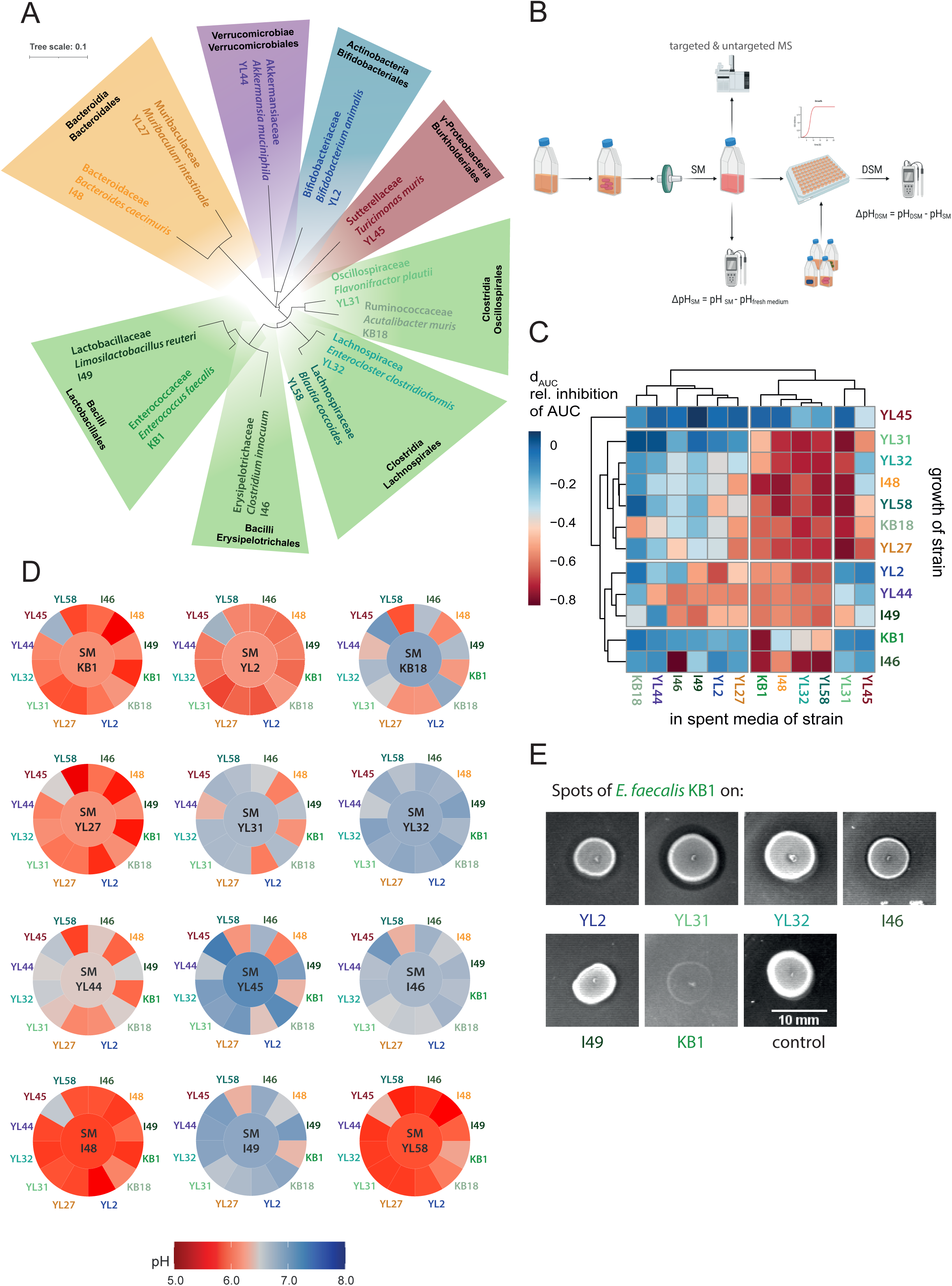
Growth analysis of OMM^12^ strains in spent media experiments. (**A**) Phylogenetic tree for bacteria of the OMM^12^ consortium based on the individual 16S rRNA genes. The consortium represents the five major phyla of the murine gastrointestinal tract: Firmicutes (green), Bacteroidetes (orange), Verrucomicrobia (purple), Actinobacteria (blue) and Proteobacteria (red). (**B**) Flowchart depicting spent culture medium (SM) preparation by growing bacterial monocultures in fresh AF medium for 20h. Culture supernatants were sterile-filtered, samples for pH measurements and MS were collected, and the SM was used as culture medium for the growth of all respective other strains. After growth of the individual strains in the specific SM, pH of the double spent medium (DSM) was determined. Differences in pH were then analyzed by calculating the corresponding ΔpH_SM_ and ΔpH_DSM_. (**C**) Monoculture growth in SM resulted in mostly decreased area under the growth curve (AUC) values in comparison to fresh AF medium, which was analyzed by calculating the inhibition factor dAUC. dAUC was calculated from the mean AUC of three independent experiments relative to the mean AUC in fresh medium. (**D**) The mean pH of all SM (center of circles) and DSM (outer tiles) after growth of the individual strains in fresh medium and the respective SM was determined from three independent experiments. (**E**) Spot assays to determine production of antibacterial agents. All bacterial strains of the OMM^12^ consortium were spotted onto a bacterial lawn of all the respective other strains. Inhibition zones were observed for *B. animalis* YL2, *F. plautii* YL31, *E. clostridioformis* YL32, *C. innocuum* I46 and *L. reuteri* I49 when *E. faecalis* KB1 was spotted. No inhibition zone was seen for *E. faecalis* KB1 on itself. AF medium with *E. faecalis* KB1 spotted is shown as control.

## Results

### Probing directional interactions of OMM^12^ strains using spent culture media

To characterize directional interactions of the OMM^12^ consortium members, we chose an *in vitro* approach to explore how the bacterial strains alter their chemical environment by growth to late stationary phase.

Growth of the individual monocultures in a rich culture medium (AF medium, Methods, **Tab. S1, Tab. S2)** was monitored over time (**Fig. S1**) and growth rates (**Tab. S3**) were determined. Strains were grouped by growth rate (GR) into fast growing strains (GR > 1.5 h^-1^, *E. faecalis* KB1, *B. animalis* YL2, *C. innocuum* I46 and *B. coccoides* YL58), strains with intermediate growth rate (GR > 1 h^-1^, *M. intestinale* YL27, *F. plautii* YL31, *E. clostridioformis* YL32, *B. caecimuris* I48 and *L.reuteri* I49) and slow growing strains (GR < 1 h^-1^, *A. muris* KB18, *A. muciniphila* YL44 and *T. muris* YL45). All strains reached late stationary phase within 20 h of growth. To probe overlap in substrate requirements and interactions between the individual OMM^12^ members mediated by waste products or bacteriocins, sterile spent culture medium (SM) after growth to late stationary phase of all strains was obtained. Each OMM^12^ strain was cultured in the SM of the other community members and their own SM and growth rate, the area under the growth curve (AUC) and the pH were determined (**Fig. 1B**, **Fig. S2)**.

A normalized inhibition factor (dAUC) was determined by the AUC in SM relative to the AUC in fresh AF medium 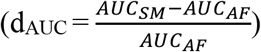 to quantify the influence of the different SM on the growth of the individual OMM^12^ strains (**Fig. 1C**). Ten of the twelve SM were found to strongly decrease (dAUC < - 0.5) the growth of at least one other strain of the consortium. Only the SM of strains *A. muris* KB18 and *A. muciniphila* YL44 were found to strongly inhibit growth of just the strains themselves. Corresponding to decreased AUC values in SM, growth rates were found to be lower as well, resulting in linear correlation of AUC and growth rates (**Fig. S3**, R > 0.5, p < 0.05 for all strains). The SM of four strains, *E. faecalis* KB1, *B. coccoides* YL58, *E. clostridioformis* YL32 and *B. caecimuris* I48, were found to strongly inhibit (dAUC < - 0.5) the growth of nine other strains each (**Fig. 1C**). Notably, growth of *E. faecalis* KB1 itself was only strongly reduced in its own SM, while it was able to grow in other strains’ SM. *T. muris* YL45 was the only strain not showing clear growth inhibition in any of the SM while its SM strongly decreased growth (dAUC < - 0.5) of three other strains, *A. muris* KB18, *M. intestinale* YL27 and *F. plautii* YL31.

### Individual pH profiles as indicators for niche modification

The pH of the culture medium after growth to stationary phase can be used as a measure for the extent of strain specific environmental modification (11) and may partly explain inhibition of bacterial growth in a SM. Therefore, we determined the pH of the individual SM before and after (double spent media; DSM) growth of all OMM^12^ strains (**Fig. 1B**, Methods). From these values, we defined the ΔpH for every strain after growth in fresh medium (ΔpH_SM_) and in all SM (ΔpH_DSM_) by analyzing the strength (difference of pH values) and direction (more acidic or more alkaline) of the pH change (**Fig. 1D**). After growth in fresh AF medium with neutral pH of 7.0, the OMM^12^ strains showed different degrees of ΔpH_SM_. While *E. faecalis* KB1, *B. animalis* YL2, *M. intestinale* YL27, *B. caecimuris* I48 and *B. coccoides* YL58 distinctly acidified the medium (pH_SM_ < 6.2), the growth of the other strains resulted in either slightly more alkaline or nearly neutral medium. Correlating inhibition of growth in a SM (dAUC) with the mean pH of the individual SM for each strain revealed that growth inhibition did not directly correlate with the pH. Only strains *B. animalis* YL2, *A. muciniphila* YL44 and *B. caecimuris* I48 showed a significant negative correlation (R < - 0.5, p < 0.05) between growth inhibition and pH (**Fig. S4**) with stronger inhibition in more acidic pH ranges.

Most interestingly, many strains did not show the same magnitude or direction of alteration in pH when grown in SM of another strain (ΔpH_DSM_) compared to growth in fresh culture medium (ΔpH_SM_). This indicates an altered metabolic behavior of some strains in specific SM environments that differs from metabolic behavior in fresh AF medium (**Fig. S5**, Supplemental Text A).

### Production of antibacterial compounds by *E. faecalis* KB1

Growth inhibition in SM (**Fig. 1C**) can further be explained by the production of antimicrobial compounds. To test the production of antimicrobial compounds by the OMM^12^ strains, we used a phenotyping approach and performed spot assays on agar plates (**Fig. 1E**). Inhibition zones were only seen in case of *E. faecalis* KB1, which produced one or several compounds active against *B. animalis* YL2, *F. plautii* YL31, *E. clostridioformis* YL32, *C. innocuum* I46 and *L. reuteri* I49. Genome analysis revealed that the strain encodes genes for the production of several bacteriocins (Supplemental Text B), including enterocin L50, an enterococcal leaderless bacteriocin with broad target range among Gram-positive bacteria (33). All other strain pairs did not show signs of growth inhibition by compound excretion under these conditions, despite the presence of genes for lanthibiotic production in the genome of *B. coccoides* YL58 (determined by antiSMASH) (34). Although expression of antimicrobial molecules may be induced by specific environmental triggers which are absent in the monoculture *in vitro* setting, we concluded that interference competition may only play a role in a subset of pair-wise interactions in AF medium involving *E. faecalis* KB1.

### Substrate depletion profiles correlate with growth inhibition in SM

As pH and antimicrobial compounds only partly explained inhibition of growth in SM, we set out to gain more insights into the individual metabolic profiles in our *in vitro* setting. Therefore, triplicate samples of fresh AF medium and SM were analyzed by a mass spectrometry-based untargeted metabolomics approach (TripleTOF, Methods). Combining positive and negative ionization mode, 3092 metabolomic features were detected in total (Methods). From these, 2387 (77.20 %) were significantly altered (t-test, p value < 0.05) by at least one of the twelve strains (**Fig. S6**). Hierarchical clustering of the metabolomic feature depletion profiles (i.e. substrates used by the bacteria; **Fig. 2A**) reflects the phylogenetic relationship between the strains (**Fig. 1A**). Correlating the phylogenetic distance between the individual strains with the number of shared depleted metabolomic features in AF medium (**Fig. S7**) showed that phylogenetically similar strains of the consortium have a higher substrate overlap than phylogenetically distant strains (R = - 0.29, p = 0.017). The total number of metabolomic features that are depleted from AF medium greatly varies for the different strains, ranging from over 600 depleted features for *M. intestinale* YL27 to only 42 for *A. muciniphila* YL44 (**Fig. 2B**). The strain specific profiles of depleted metabolomic features were compared pairwise and the number of overlapping features was determined (**Fig. 2C**). Phylogenetically related strains like *E. clostridioformis* YL32 and *B. coccoides* YL58 or *M. intestinale* YL27 and *B. caecimuris* I48 share over 50% of depleted metabolic features each, suggesting a strong substrate overlap in AF medium. Visualizing the extend of overlap between substrate depletion profiles reveals that Bacteroidales, Clostridia and Bacilli strains of the consortium dominate with the highest number of commonly depleted substrates in AF medium (**Fig. 2D**).

**Figure 2:**
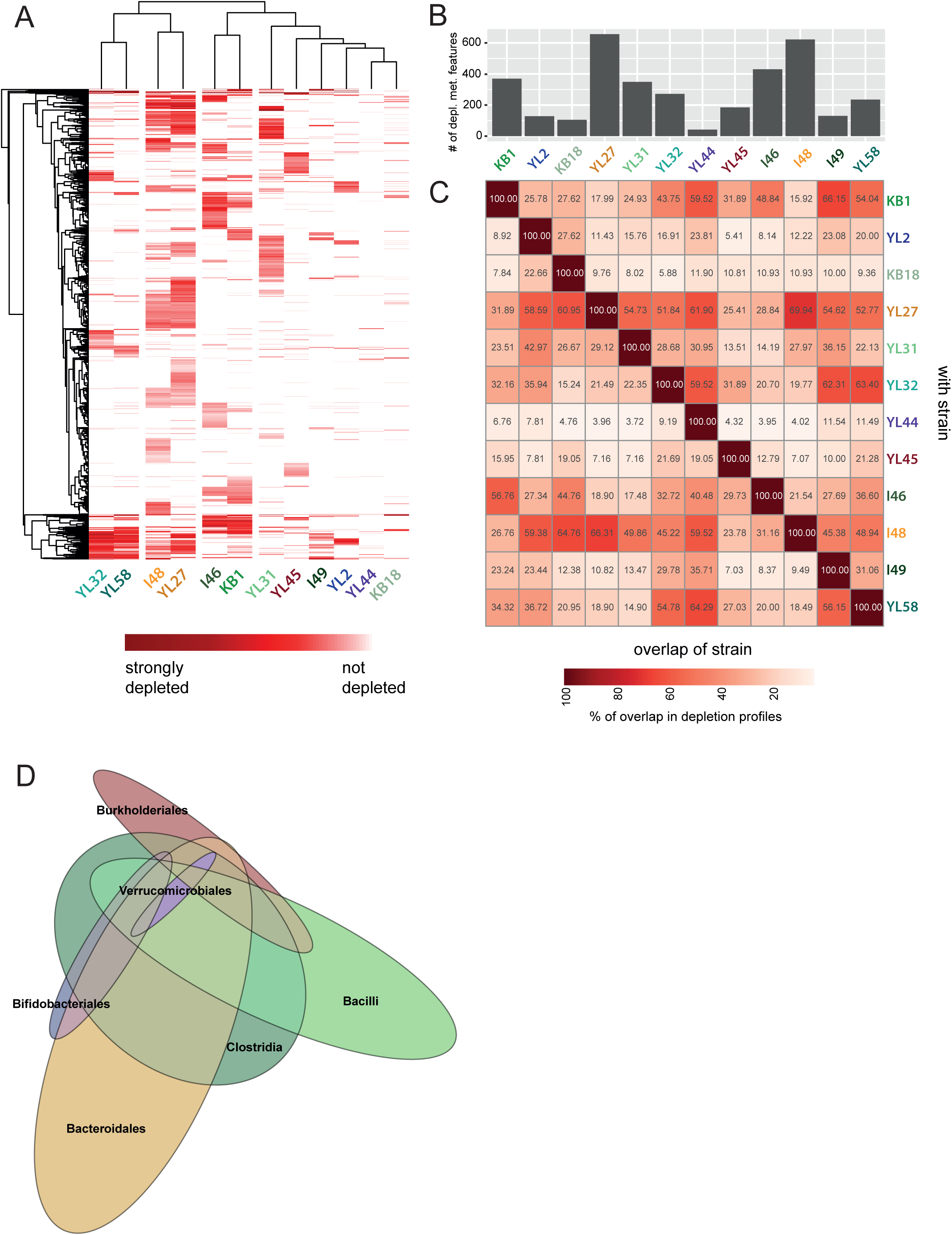
Overlap of substrate depletion profiles between individual OMM^12^ strains. (**A**) Depletion profiles of substrates after bacterial growth to stationary phase in AF medium were determined by untargeted MS from three independent experiments. All metabolomic features (rows) that significantly decreased (p < 0.05 compared to fresh media) compared to fresh medium for at least one of the twelve strains are shown in red. Dark-red indicates strong depletion, while white indicates no depletion of the metabolomics feature. Hierarchical clustering of strain specific profiles as well as metabolomic features reveal profile similarities between phylogenetically similar strains. (**B)** Bar plot showing the total number of significantly (p < 0.05 compared to fresh media) depleted metabolomic features in AF medium for the individual strains. (**C**) Pairwise overlap in depletion profiles in percent of the total number of individually depleted metabolomic features. (**D**) Euler diagram depicting total number of depleted metabolomic features and overlap within the consortium. Size of the ellipse denotes number of depleted features, size of overlap between ellipses denotes number of features that are shared when comparing the individual profiles.

Correlating the growth inhibition in SM (d_AUC_) with the pairwise overlap in depletion profiles (**Fig. 2C**) revealed that a larger overlap is correlated with a stronger growth inhibition in the corresponding SM (R = - 0.46, p = 3.1E-08, **Fig. S8**). This is illustrated by *A. muciniphila* YL44, which used only a low numbers of substrates from the AF medium (**Fig. 2B**) and the SM of which had only little effect on the growth of the other strains of the consortium (**Fig. 1C**). On the other hand, the strain’s growth itself was strongly reduced in the SM of most other consortium members (**Fig. 1C, Fig. S2**), which depleted a large spectrum of metabolomic features including those used by *A. muciniphila* YL44 (**Fig. 2C**).

### Genome-informed metabolic potential of the OMM^12^ consortium

To be able to infer metabolic interactions between the individual consortium members, a reference dataset giving insight into the metabolic potential of the OMM^12^ based on genetic information was generated. We screened the genomes of the twelve strains for key enzymes of central carbon metabolism (e.g., fermentation pathways, respiration and amino acid metabolism) (**Fig. 3A**), as well as for transporters (ABC-transporters and PTS-systems) for carbohydrates and amino acids (**Fig. S9, SI data table**). Hierarchical clustering of the genome-informed metabolic potential (**Fig. 3A**) reflected phylogenetic relationships in several instances, e.g., the Lachnospirales strains of the consortium *E. clostridioformis* YL32 and *B. coccoides* YL58, as well as the Oscillospirales strains *F. plautii* YL31 and *A. muris* KB18 were found to cluster closely together. Of note, the metabolic potential of *T. muris* YL45 (Sutterellaceae) was very distinct, clustering differently from all other strains. Generally, high diversity of central and fermentation pathways was found among the consortium members. Moreover, enzymes for the degradation of monosaccharides (e.g., arabinose, xylose and ribose) and amino acids (e.g., methionine and glutamine) are highly prevalent among consortium members. Phosphotransferase systems were especially prevalent among strains *E. faecalis* KB1, *E. clostridioformis* YL32 and *C. innocuum* I46, while ABC transporters for carbohydrates and amino acids were more distributed among all consortium members (**Fig. S9**).

**Figure 3:**
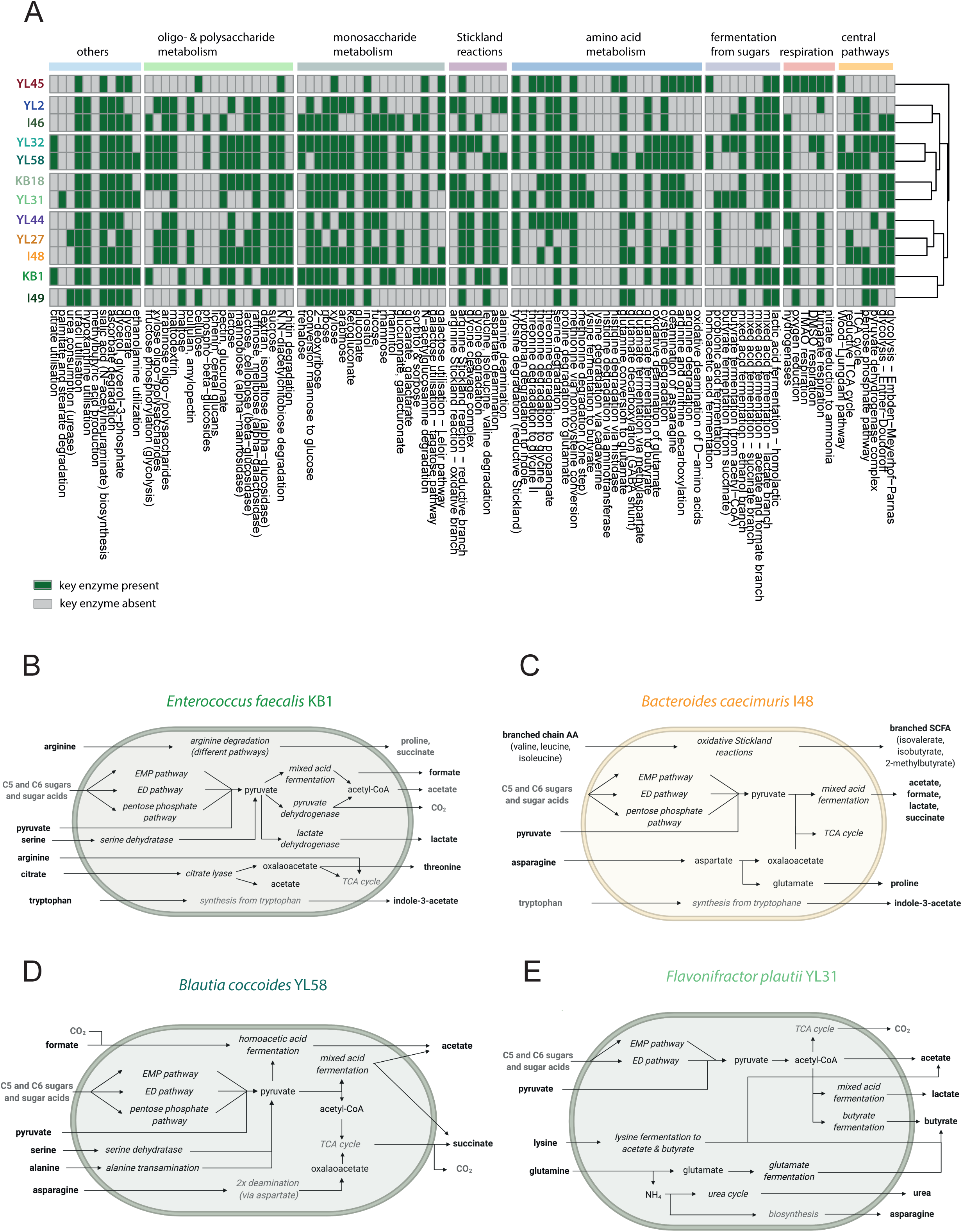
Metabolic potential of the OMM^12^ strains. (**A**) OMM^12^ genomes were screened for a hand-curated set of key enzymes to determine the strains’ potential to use a diverse range specific substrates, metabolic pathways and release fermentation end products. A potential substrate and pathway utilization was considered positive (green) if one of the associated KO’s was found in the respective genome (**SI data table**). Consequently, positive hits do not indicate completeness of the pathway. If none of the associated KO’s was found in the respective genome, the potential substrate and pathway utilization was considered negative (grey). Metabolites and pathways were sorted by functional groups. (**B**) By combining metabolomics data (MS, **Fig. S10, S11**) with genome-based information on the presence of key enzymes, broad-scale draft metabolic models of the individual OMM^12^ strains were generated (supplementary text C, **Fig. S12**). Here, the models for strains *E. faecalis* KB1, (**C**) *B. caecimuris* I48, (**D**) *B. coccoides* L58 and (**E**) *F. plautii* YL31 are shown. Models of remaining strains of the consortium are shown in **Fig. S12**. Experimentally confirmed substrates, products or enzymes are shown in black. Hypothetical substrates, products or enzymes are shown in grey.

### Metabolite production and fermentation pathways of the OMM^12^ strains in AF medium

To gain insights into the metabolites and fermentation products produced and consumed by the individual strains of the consortium in the given *in vitro* conditions, SM were analyzed using different mass spectrometry approaches (Methods, **Fig. S10, Fig. S11**). Combining experimentally obtained insights with genome-based information on the presence of key enzymes enabled the generation of broad-scale draft metabolic models of the individual OMM^12^ community members (**Fig. 3B-E**, **Fig. S12,** Supplemental Text C).

To confirm that fermentation pathways identified by genomics were active under *in vitro* conditions, short chain fatty acid (SCFA) production and consumption was analyzed (**Fig. S10A**). As observed for the SM metabolic profiles (**Fig. S6**), hierarchical clustering revealed that closely related bacteria showed similar SCFA production and consumption profiles. Both Bacteroidetes strains produced acetic acid, succinic acid as well as branched-chain fatty acids. Both Lachnospiraceae strains generated high amounts of acetic acid. Butyric acid is produced by strains *F. plautii* YL31 and *C. innocuum* I46, the latter also being the only strain of the consortium excreting valeric acid and hexanoic acid. Of note, *F. plautii* YL31 also consumed lysine, indicating the ability to produce butyric acid from lysine, which was supported by the presence of gene coding for lysine aminomutase (EC 5.4.3.2 and EC 5.4.3.3) as well as two of the following genes encoding key enzymes in the pathway: L-erythro-3.5-diaminohexyanoate dehydrogenase (EC 1.4.11) and 3-keto-5-aminohexanoate cleavage enzyme (EC 2.3.1.247).

Formic acid was produced by several strains and consumed by *T. muris* YL45 and *B. coccoides* YL58, indicating the ability of formic acid/ H2 oxidation. *T. muris* YL45 and *B. coccoides* YL58 both encode genes for a CO dehydrogenase/acetyl-CoA-synthase (EC 1.2.7.4 and EC 2.3.1.169), the key enzyme of the Wood-Ljungdahl pathway (reductive acetyl-CoA pathway). Formic acid can be processed via this pathway to acetyl-CoA. As another prominent example of bacterial fermentation, lactate production was confirmed (**Fig. S10B)** for *E. faecalis* KB1, *B. animalis* YL2, *F. plautii* YL31, *A. muris* YL45, *C. innocuum* I46 and *B. caecimuris*, all of which harbor genes coding for the enzyme lactate dehydrogenase (EC 1.1.1.27 and EC 1.1.1.28).

By quantifying amino acid levels we could show that Bacteroidetes and Lachnospiraceae strains exhibited similar amino acid depletion and production profiles. In SM of strains *M. intestinale* YL27 and *B. caecimuris* I48, elevated levels of a diverse range of amino acids including glutamic acid, histidine, methionine, proline and phenylalanine were detected. Lachnospiraceae strains showed increased levels of isoleucine, tryptophan and valine, while alanine was especially depleted by *B. coccoides* YL58. Other strains of the consortium showed specific depletion of single amino acids, e.g., *F. plautii* YL31 strongly depleted lysine and glutamic acid, while *E. faecalis* KB1 depleted serine.

### Growth of OMM^12^ strains in pairwise co-culture

Next, we performed a set of experiments to characterize strain-strain interactions in the dynamic community-dependent context. We first analyzed direct competition of all strains in pair-wise co-cultures over the course of 72h, with serial dilutions every 24h. While growth was monitored continuously by OD 600nm, samples for pH measurements and qPCR analysis were taken every 24h. The growth curves of most co-cultures, as well as supernatant pH differed from the corresponding strain specific characteristics observed in monoculture (**Fig. S13, Fig. 4A**). These differences reflect co-culture dynamics, as can be seen from change in relative abundances over time (**Fig. 4B, C**).

**Figure 4:**
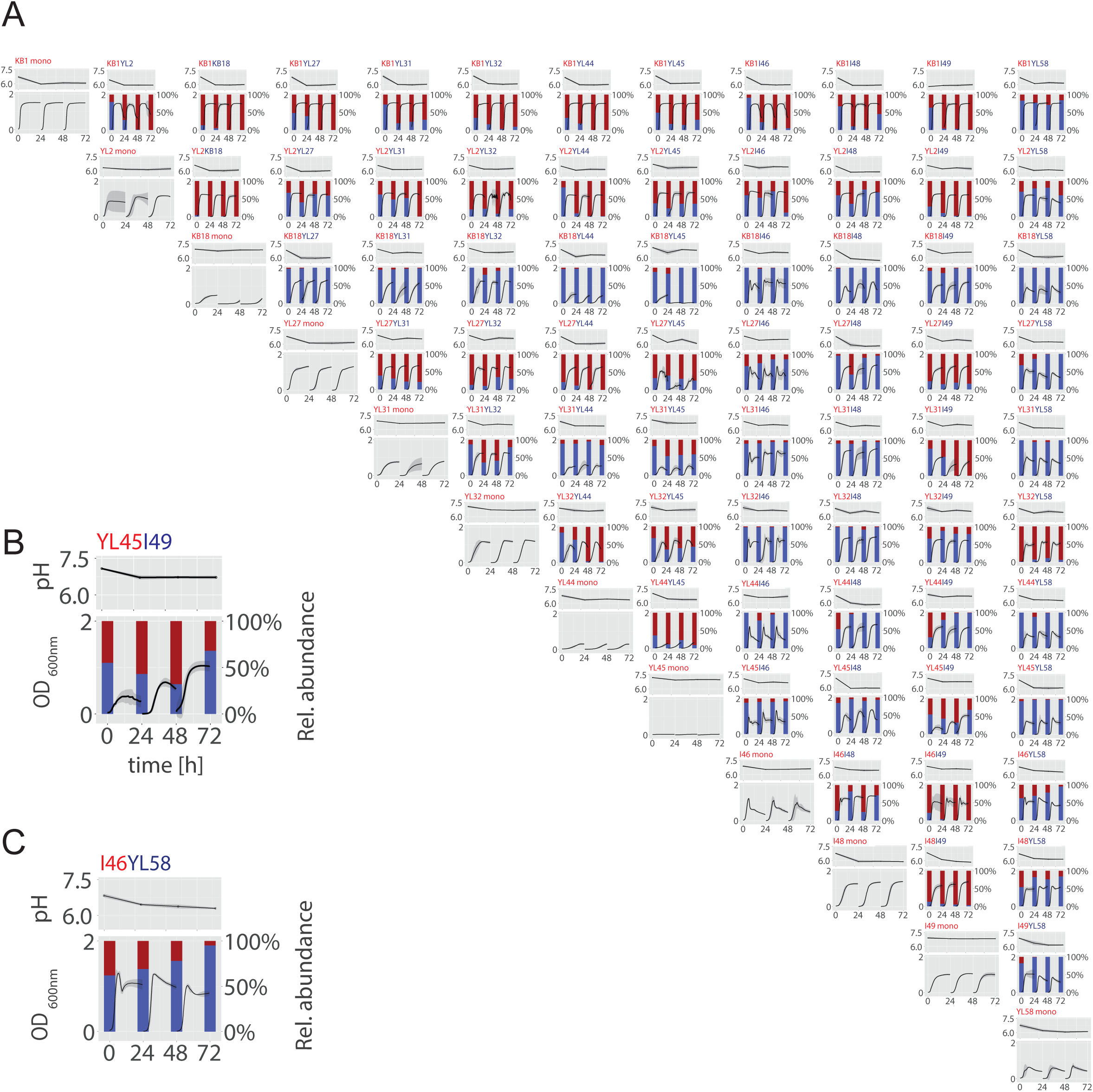
Pairwise cultures of the OMM^12^ strains. (**A**) OMM^12^ pairwise strain combinations (12 monocultures, 66 co-cultures) were cultured in a 1:1 ratio in fresh AF medium over the course of 72 hours and growth, pH and relative abundance was monitored over time in three independent experiments. Growth at OD_600nm_ and pH is shown as mean with the corresponding standard deviation (grey), relative abundance over time is shown exemplary for one of the three experiments. Examples of how growth curves develop with changing relative abundances is shown for the co-culture of *T. muris* YL45 and *L. reuteri* I49 (**B**). Starting with a OD_600nm_ ratio of approximately 1:1, final mean OD_600nm_ values after the first two turnovers are low, corresponding to YL45 dominating the co-culture. After 48h, *L. reuteri* I49 resumes growth and final OD values increase. With *L. reuteri* I49 dominating the community in the end, the growth curve as well resembles *L. reuteri* I49 monoculture growth. Similarly, pH values reflect changes in co-culture structure, as can be observed e.g. in the co-culture of *C. innoccum* I46 and *B. coccoides* YL58 (**C**). In monoculture, *C. innocuum* I46 does not strongly acidify its environment, while YL58 acidifies the culture supernatant to around pH 6.0. In accordance with this observation, pH values of the co-culture supernatant only drop with increasing dominance of YL58 in the co-culture.

To identify directionality and mode of interaction between the OMM^12^ strains, we analyzed the relative changes in absolute abundance (16S rRNA gene copies) as a measure of how successful a strain can grow in co-culture relative to monoculture after 72h. The mean absolute abundance ratio was calculated for every strain in all pairwise co-cultures 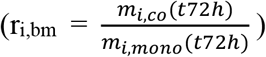 (**Fig. S14**, Methods). If absolute abundance of a strain increased significantly in co-culture relative to monoculture (r_bm_ > 1), the interaction was categorized as positive (+), if it decreased (r_bm_ < 1) the interaction was categorized as negative (-) (t-test comparing the rbm of three independent experiments, **Fig. S15**). If it did not significantly (p > 0.05) differ from that in monoculture (r_bm_ = 1), the interaction was categorized as neutral (0). By this, we created a co-culture interaction matrix (**Fig. 5A**): the vast majority of the interactions was classified as amensalistic (0/- and -/0, 46 of 66 of interactions). A smaller subset of interactions was either competitive (-/-, 7 of 66 of interactions) or neutral (0/0, 11 of 66 of interactions). No mutualistic interactions (+/+) were observed. However, one example for each, commensalism (0/+ and +/0) and predation (+/− and -/+), were identified.

**Figure 5:**
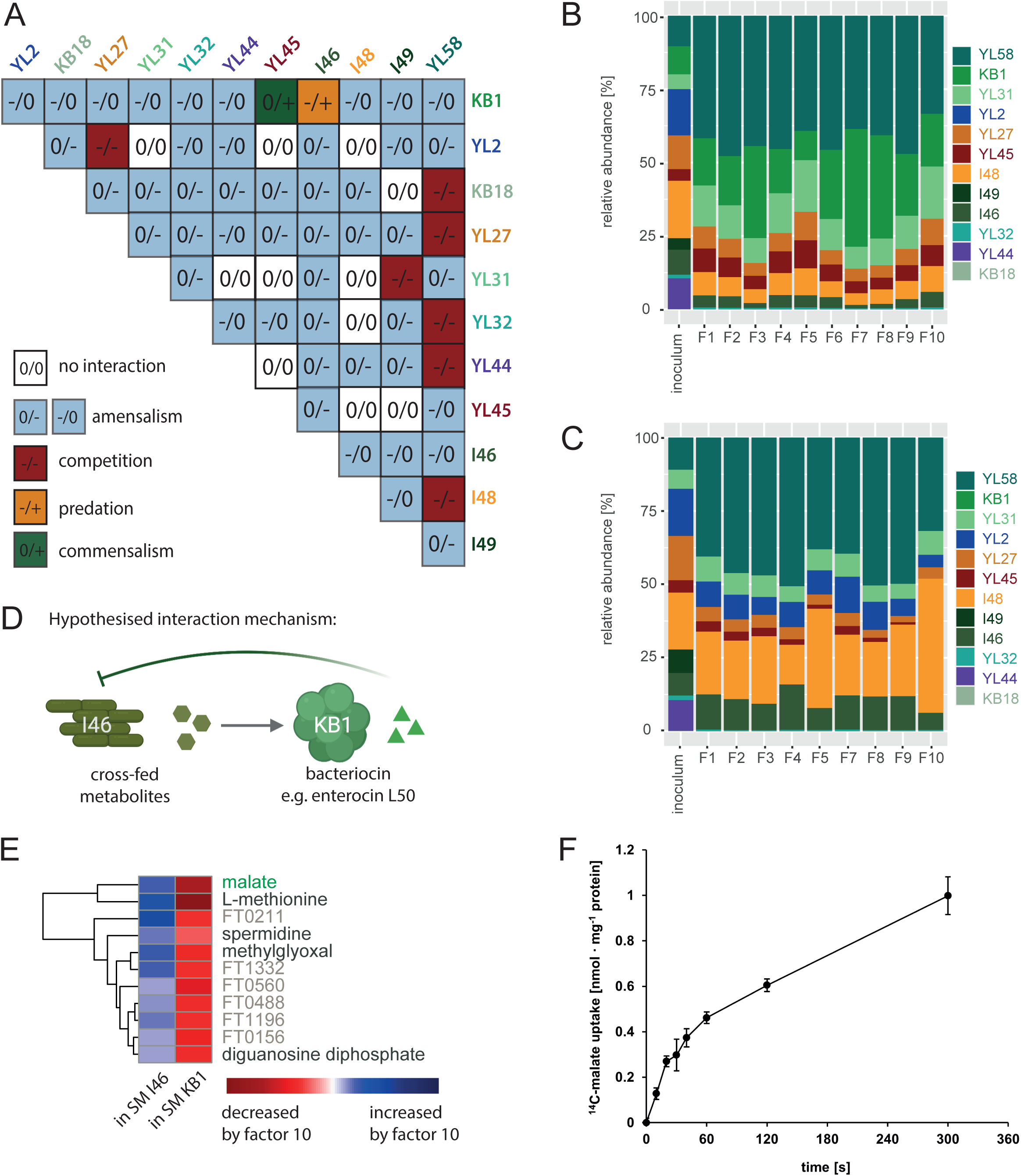
Transferring pairwise interactions to the community level. (**A**) By comparing the mean bacterial abundance from three independent experiments in co-culture to the mean abundance in monoculture, a pairwise interaction matrix was generated. Interactions where the individual abundance in co-culture significantly (t-test, p< 0.05) increased are indicated with (+), interactions where it significantly decreased are indicated with (–) and interaction where the abundance did not change in comparison to monoculture growth were indicated with (0). (**B**) Using a serial passaging batch culture setup, the OMM^12^ community composition was analyzed after ten days of serial dilutions by comparing the relative strain abundances of ten replicates (F1-F10) with the inoculum. (**C**) To study the influence of *E. faecalis* KB1 on community composition, a OMM^11^-KB1 dropout community was constructed and relative abundance at day ten was analyzed for ten replicates (F1-F10). (**D**) Based on SM experiments, pairwise co-culture and community experiments a predation interaction mechanism between strains *C. innocuum* I46 and *E. faecalis* KB1 is proposed. We hypothesize, that KB1 produces a bacteriocin inhibiting I46, and benefits from cross-feeding on I46 derived metabolites. (**E**) Potentially cross-fed metabolites were determined by comparing SM profiles (determined by untargeted MS) of KB1 and I46 for metabolites that are strongly produced by I46 and consumed by KB1. Verified annotations are shown in green, potential annotations are shown in black and not annotated compounds are shown in grey as the corresponding feature identification numbers. (**F**) Time course of malate uptake by whole cells of *E. faecalis* KB1. Rates of ^14^C-malate uptake were measured at a final malate concentration of 10 μM at 18° C. Standard deviations are estimated from three biological replicates.

The extent to which the individual strains altered the growth of other community members in the co-culture differed distinctly. While *E. faecalis* KB1 and *C. innocuum* I46 lead to nine negative co-culture outcomes each, *A. muciniphila* YL44 and *A. muris* KB18 only impaired growth of one and zero strains, respectively. Simultaneously, both strains are negatively influenced in most co-cultures, with a significantly decreased absolute abundance in ten and eight co-cultures, respectively. Notably, *B. coccoides* YL58 is involved in five of seven competitive interactions of the consortium. These observations are in line with the outcomes observed in SM experiments, as strongly negative co-culture outcome correspond to a strong inhibition of a strain in the respective SM (**Fig. S16**).

### Community structure of the OMM^12^ consortium

Next, we set out to investigate if interactions found in co-cultures are transferrable to the strains’ behavior in the complete OMM^12^ community. To this end, all twelve OMM strains were simultaneously co-cultured in AF medium and were serially diluted 1:100 every 24h into fresh AF medium. Relative abundance of all strains after 72h and 10 days compared to the inoculum was determined by qPCR for ten replicates each in two independent experiments from different inocula (**Fig. S17, Fig. S18, Fig. 5B**).

While each of the OMM^12^ members except *E. faecalis* KB1 was outcompeted to a very low relative abundance in at least one pairwise culture (**Fig. 5A**), the majority (10 out of 12) of the consortium members were able to coexist in the complex community over the course of 72h (**Fig. S17A**) and up to 10 days. (**Fig. 5B, Fig. S17B**). Replicate communities showed reproducible community structure, even when different inocula were used (**Fig. 5B, Fig. S17B**) and especially when compared on the order and phylum level (**Fig. S18**). Communities were dominated by *B. coccoides* YL58 and *E. faecalis* KB1, together making up > 50% of the relative abundance, which corresponds to their dominant role in SM and co-culture experiments (**Fig. 1C, 5A**). While strains *B. animalis* YL2 and *L. reuteri* I49 were not detectable at 72h and 10 days in all replicates, *A. muris* KB18 was found in only few of the communities at 10 days (relative abundance < 1%).

### *E. faecalis* KB1 strongly impacts overall community composition

To understand how interactions observed on the pairwise level transfer to the community context, we derived pairwise correlations from the relative abundance data of all communities at day 10 derived from two independent experiments (**Fig. S19**). The strong negative influence of *E. faecalis* KB1 on most other strains observed in pairwise co-cultures applied in the community context as well. High relative abundance of *E. faecalis* KB1 linearly correlated with decreased relative abundance of *C. innocuum* I46 (R = - 0.87; p < 0.05), supporting the hypothesis of a predatory interaction between *E. faecalis* KB1 and *C. innocuum* I46 as observed in co-culture experiments (**Fig. 5A, D**). In order to identify potentially cross-fed metabolites of *C. innocuum* I46 to *E. faecalis* KB1, we mined metabolomic data of SM for features enriched in *C. innocuum* I46 and depleted by *E. faecalis* KB1. Thereby, we identified several compounds including malate, L-methionine, spermidine and methylglyoxal (**Fig. 5E**). To experimentally support the idea of cross-feeding, we exemplarily tested uptake of ^14^C-malate into intact cells of *E. faecalis* KB1. To slow the metabolization of this metabolite, all assays were performed at 18° C. We found a very fast linear uptake of ^14^C-malate by *E. faecalis* KB1 within the first 60 s, which could explain why malate utilization confers a growth advantage to this strain **(Fig. 5F)**.

Finally, we were interested in how the absence of *E. faecalis* KB1 would affect the overall community structure. We generated a ‘dropout’ community including all strains of the OMM^12^ consortium except *E. faecalis* KB1 (OMM^11^-KB1). Compositional analysis revealed increased relative abundance of *C. innocuum* I46 and *B. animalis* YL2 in the OMM^11^-KB1 compared to the full OMM^12^ community (**Fig. 5C**). In addition, the absolute abundances of strains *B. animalis* YL2, *C. innocuum* I46 and *B. caecimuris* I48 were found to increase significantly (t-test, p < 0.05) in the absence of *E. faecalis* KB1 (**Fig. S20**). While the increase in abundance of *B. animalis* YL2 and *C. innocuum* I46 may be explained by absent enterocin production by *E. faecalis* KB1, the increased abundance of *B. caecimuris* I48 was unexpected. Further, the abundance of *F. plautii* YL31, *E. clostridioformis* YL32, *A. muciniphila* YL44 and *T. muris* YL45 was found to decrease in the absence of *E. faecalis* KB1. This indicates either direct positive effects of *E. faecalis* KB1 on these strains or indirect effects that occur through the overall shift in OMM^11^-KB1 community composition compared to the OMM^12^ consortium.

## Discussion

A central challenge in gut microbiome research is to understand how interactions between the individual microorganisms affect community-level structure and related functions. Bottom-up approaches involving synthetic communities are valuable tools to study these interactions, as they allow to reduce complexity and to enable strain-specific manipulation. Using an *in vitro* approach, we focused on characteristics and interactions of the OMM^12^ community and combined monoculture, pairwise and community cultivation of the strains with genome and metabolomics analysis of their SM. Thereby we reveal that the OMM^12^ community interaction network is shaped by exploitative and interference competition. In particular, *E. faecalis* KB1, a low-abundant member of the mammalian gut microbiota, was identified as important driver of *in vitro* community composition by directly or indirectly altering the abundance of several other consortium members. We provide draft metabolic models of the individual OMM^12^ strains, which will be a valuable tool for mechanistic studies using this synthetic community.

Exploitative (i.e. substrate) competition plays a major role in shaping intestinal bacterial communities (35). Understanding the underlying principles of how bacteria compete for available nutrients is essential to predict and control community composition. We found that phylogenetically similar strains showed a higher substrate overlap (**Fig. S7**), which is in accordance with previous studies demonstrating that phylogeny reflects metabolic capabilities of bacteria (36, 37). Furthermore, overlap in substrate depletion profiles was correlated with growth inhibition in the respective SM (**Fig. S8**). This clearly indicates strong exploitative competition between individual OMM^12^ strains. In particular *B. caecimuris* I48, *E. faecalis* KB1, *E. clostridioforme* YL32 and *B. coccoides* YL58 were found to consume a high number of substrates (>200), while their SM inhibited growth of the majority of the other community members (**Fig. 1C**). Of note, *M. intestinale* YL27 and *C. innocuum* I46, also consumed over 200 substrates each, but inhibited few other strains. This demonstrates that substrate overlap cannot simply predict inhibition in all cases and other mechanisms (i.e. waste product inhibition) play a role in specific cases.

Besides substrate competition, a strain’s ability to acidify its environment or release an inhibitory factor (e.g. waste products, bacteriocins) can determine if another species can grow in the exhausted medium or not. Several strains, including *B. caecimuris* I48, *B. coccoides* YL58, *E. faecalis* KB1 and *C. inocuum* I46 acidified the medium during growth in monoculture. However, only for few species, *A. muciniphila* YL44, *B. caecimuris* I48 and *B. animalis* YL2, acidic pH correlated with reduced growth (**Fig. S4**). Moreover, the pH in the full OMM^12^ community, where most of the strains coexisted, was also acidic (pH of 6.2), suggesting that pH modification does not play a major role in driving community composition *in vitro*. Interference competition by bacteriocins is widespread among gut bacterial communities (38). We found that *E. faecalis* KB1 produces at least one antimicrobial compound that shows activity against five of the Gram-positive OMM^12^ strains (*B. animalis* YL2, *E. clostridioformis* YL32, *F. plautii* YL31, *C. innocuum* I46 and *L. reuterii* I49) (**Fig. 1E**). *E. faecalis* harbors genes coding for at least two different enterocins (enterocin L50A/L50B and enterocin O16). Therefore, we hypothesize that some of the inhibitory effects of *E. faecalis* KB1 on those strains can be attributed to enterocin-mediated killing.

*E. faecalis* is a prevalent but low abundant member of the undisturbed human and animal microbiota. Following antibiotic therapy, the bacterium can dominate the gut and cause blood-stream infection in immunocompromised individuals (39). Understanding how *E. faecalis* out-competes/overgrows other gut microorganisms is important in order to intervene with *E. faecalis* domination in the gut. Besides enterocin-mediated killing we found that metabolite cross-feeding seems to contribute to the interaction of *E. faecalis* KB1 with *C. innocuum* I46 (**Fig. 5D; Fig. S10**). Based on metabolic profile mining, we hypothesize, that *E. faecalis* KB1 consumes malate, methionine, arginine and serine among other metabolites in co-culture with *C. innocuum* I46 (**Fig. 5E, Fig. S11**). Interestingly, a previous study (40) reported that glucose-malate co-metabolism increases growth of *E. faecalis* over glucose consumption alone. In connection with fast uptake rate of ^14^C-malate by *E. faecalis* KB1 (**Fig. 5F**), this suggests that malate cross-feeding may also contribute to *E. faecalis* KB1 gain in absolute abundance in coculture with *C. inoccuum* I46.

Using batch culture experiments we were able to investigate assembly and dynamics of full OMM^12^ and OMM^11^-KB1 dropout communities *in vitro*. A significant increase in *B. animalis* YL2 and *C. innocuum* I46 in the community lacking *E. faecalis* KB1 suggested that enterocin-mediated killing also shapes the more complex community (**Fig. S20**). Notably, in the full OMM^12^ community, ten of the twelve strains co-existed over ten days. This was remarkable given the high number of negative pairwise interactions in SM and co-culture experiments. Differences between the behavior of strains in pairwise versus complex communities point at higher-order ecological interactions that emerge in the community context. As previously shown in other studies, the underlying mechanisms may involve metabolic flexibility or mixed substrate utilization of the strains in the presence of competitors (41), metabolite cross-feeding and lack of waste-product inhibition and overall change in pH (11) or an excess of provided substrates in the medium.

Following up, it will be important to assess, if the *in vitro* findings can be translated to the mouse model. Several differences between *in vitro* and *in vivo* conditions were noted. While the *in vitro* community is dominated by *B. coccoides* YL58 and *E. faecalis* KB1, mouse communities are dominated by *B. caecimuris* I48 and *A. muciniphila* YL44 (**Fig. S21**)(26). Enrichment of amino acids and glucose in the used culture medium may favor growth of *B. coccoides* YL58 and *E. faecalis* KB1 *in vitro* at the expense of bacteria specialized on utilization of mucin and other complex carbohydrates (42, 43). To this end, it will be worthwhile to modify the *in vitro* conditions to more closely recapitulate the chemical landscape and spatial structure of the gut in future experiments.

Concluding, our study presents a comprehensive *in vitro* investigation of strain-strain interactions between members of a widely used synthetic intestinal bacterial community. Characterization of the metabolic profile of individual strains of the consortium as well as analyzing their metabolism and community assembly in co-culture revealed *E. faecalis* KB1 and *B. coccoides* YL58 to be important drivers of community composition. Drawing on this detailed understanding of *in vitro* behavior, our results will enable to employ this model for mechanistic *in vivo* studies. This step-wise approach may ultimately allow accurate description of interaction dynamics of *in vivo* gut microbial communities and pave the way for targeted manipulation of the microbiome to promote human health. In particular, extending the approach of dropout communities lacking specific strains could help to elucidate the role of individual players in community functions like dietary breakdown, metabolite production and colonization resistance and to identify general principles of how bacterial interaction networks and the corresponding emergence of higher order interactions shape microbiome function. This will enable the design of therapeutic interventions to control microbial community functions by advanced microbiome engineering.

## Methods

### Generation of a 16S gene based phylogenetic tree

The genomes of the twelve strains of the OMM^12^ consortium (27) were accessed via DDBJ/ENA/GenBank using the following accession numbers: CP022712.1, NHMR02000001-NHMR02000002, CP021422.1, CP021421.1, NHMQ01000001-NHMQ01000005, NHTR01000001-NHTR01000016, CP021420.1, NHMP01000001-NHMP01000020, CP022722.1, NHMU01000001-NHMU01000019, NHMT01000001-NHMT01000003, CP022713.1 and annotated using Prokka (default settings) (44). The 16S rRNA sequences of all strains were obtained. These rRNA FASTA sequences were uploaded to the SINA Aligner v1.2.11 (45) to align these sequences with minimum 95% identity against the SILVA database. By this, a phylogenetic tree based on RAxML (46), GTR Model and Gamma rate model for likelihood was reconstructed. Sequences with less than 90% identity were rejected. The obtained tree was rooted using *midpoint.root()* in the phytools package (47) in R and visualized using iTOL online (48).

### Genome annotation for predicting genome-informed metabolic potential

The genomes (accessed via DDBJ/ENA/GenBank as stated above) of the twelve strains of the OMM^12^ consortium were annotated using prodigal (version V2.6.3, default settings) (49) and KEGG orthologies (KO) for the protein-coding genes were obtained using the tool KOfamscan (default settings) (50). The tool provided multiple KO annotations for each gene with corresponding e-values and threshold scores. In order to get one KO annotation per gene, the annotation was considered only if a) the internal threshold score was reached (marked ‘*’ by kofamscan) or b) an evalue of > 1e-03 was reached. The remaining annotations were ignored.

### Strains and culture conditions

Bacterial cultures were prepared from frozen monoculture stocks in a 10ml culture and subculture in cell culture flasks (flask T25, Sarstedt) previous to all experiments. Cultures were incubated at 37°C without shaking under strictly anaerobic conditions (gas atmosphere 7% H2, 10% CO2, 83% N2). All experiments were carried out using AF medium (18 g.l^-1^ brain-heart infusion, 15 g.l^-1^ trypticase soy broth, 5 g.l^-1^ yeast extract, 2.5 g.l^-1^ K_2_HPO_4_, 1 mg.l^-1^ haemin, 0.5 g.l^-1^ D-glucose, 0.5 mg.l^-1^ menadione, 3% heat-inactivated fetal calf serum, 0.25 g.l^-1^ cysteine-HCl·H2O). The following strains were used in this study: *Enterococcus faecalis* KB1 (DSM32036), *Bifidobacterium animalis* YL2 (DSM26074), *Acutalibacter muris* KB18 (DSM26090), *Muribaculum intestinale* YL27 (DSM28989), *Flavonifractor plautii* YL31 (DSM26117), *Enterocloster clostridioformis* YL32 (DSM26114), *Akkermansia muciniphila* YL44 (DSM26127), *Turicimonas muris* YL45 (DSM26109), *Clostridium innocuum* I46 (DSM26113), *Bacteroides caecimuris* I48 (DSM26085), *Limosilactobacillus reuteri* I49 (DSM32035), *Blautia coccoides* YL58 (DSM26115).

### Growth measurements

Bacterial growth was measured in 96well round bottom plates (Nunc) using a GenTech Epoch2 plate reader. Inocula were prepared from a previous culture and subculture and diluted in fresh AF medium to 0.01 OD_600nn_. Absorption at wavelength 600nm was determined in a reaction volume of 100 μl in monoculture and SM experiments and 150 μl in co-culture experiments. During continuous measurements, the plate was heated inside the reader to 37°C and a 30 second double orbital shaking step was performed prior to every measurement.

### Generation of spent culture media

Bacterial cultures and subcultures were grown for 24 hours each in 10ml AF medium at 37°C under anaerobic conditions without shaking. Bacterial spent culture supernatants (SM) were generated by centrifugation of the densely grown subculture at 4°C for 20min at 5000xg and subsequent pH measurement and filter-sterilization (0.22 μm). SM samples were aliquoted and immediately frozen at −80°C. Samples were thawed under anaerobic conditions previous to growth measurements. Growth of all bacterial monocultures in the spent culture media (SM) of all respective other strains was then measured as described above. SM were inoculated with bacterial monocultures with starting OD_600nm_ 0.01. After monoculture growth of 20h in the respective SM (resulting in double spent media, DSM), pH values were determined.

### pH measurements

pH measurements of bacterial supernatants were performed using a refillable, glass double junction electrode (Orion™ PerpHecT™ ROSS™, Thermo Scientific).

### Metabolic profiling of late stationary phase bacterial supernatants

The untargeted analysis was performed using a Nexera UHPLC system (Shimadzu) coupled to a Q-TOF mass spectrometer (TripleTOF 6600, AB Sciex). Separation of the spent media was performed using a UPLC BEH Amide 2.1×100, 1.7 μm analytic column (Waters Corp.) with 400 μL/min flow rate. The mobile phase was 5 mM ammonium acetate in water (eluent A) and 5 mM ammonium acetate in acetonitrile/water (95/5, v/v) (eluent B). The gradient profile was 100% B from 0 to 1.5 min, 60% B at 8 min and 20% B at 10 min to 11.5 min and 100% B at 12 to 15 min. A volume of 5μL per sample was injected. The autosampler was cooled to 10°C and the column oven heated to 40°C. Every tenth run a quality control (QC) sample which was pooled from all samples was injected. The spent media samples were measured in a randomized order. The samples have been measured in Information Dependent Acquisition (IDA) mode. MS settings in the positive mode were as follows: Gas 1 55, Gas 2 65, Curtain gas 35, Temperature 500°C, Ion Spray Voltage 5500, declustering potential 80. The mass range of the TOF MS and MS/MS scans were 50 - 2000 *m/z* and the collision energy was ramped from 15 - 55 V. MS settings in the negative mode were as follows: Gas 1 55, Gas 2 65, Cur 35, Temperature 500°C, Ion Spray Voltage −4500, declustering potential −80. The mass range of the TOF MS and MS/MS scans were 50 - 2000 *m/z* and the collision energy was ramped from −15 - −55 V.

The “msconvert” from ProteoWizard (51) were used to convert raw files to mzXML (de-noised by centroid peaks). The bioconductor/R package xcms (52) was used for data processing and feature identification. More specifically, the matched filter algorithm was used to identify peaks (full width at half maximum set to 7.5 seconds). Then the peaks were grouped into features using the “peak density” method (52). The area under the peaks was integrated to represent the abundance of features. The retention time was adjusted based on the peak groups presented in most of the samples. To annotate possible metabolites to identified features, the exact mass and MS2 fragmentation pattern of the measured features were compared to the records in HMBD (53) and the public MS/MS database in MSDIAL (54), referred to as MS1 and MS2 annotation, respectively. The QC samples were used to control and remove the potential batch effect, t-test was used to compare the features’ intensity from spent media with fresh media.

### Targeted short chain fatty acid (SCFA) measurement

The 3-NPH method was used for the quantitation of SCFAs (55). Briefly, 40 μL of the SM and 15 μL of isotopically labeled standards (ca 50 μM) were mixed with 20 μL 120 mM EDC HCl-6% pyridine-solution and 20 μL of 200 mM 3-NPH HCL solution. After 30 min at 40°C and shaking at 1000 rpm using an Eppendorf Thermomix (Eppendorf, Hamburg, Germany), 900 μL acetonitrile/water (50/50, v/v) was added. After centrifugation at 13000 U/min for 2 min the clear supernatant was used for analysis. The same system as described above was used. The electrospray voltage was set to −4500 V, curtain gas to 35 psi, ion source gas 1 to 55, ion source gas 2 to 65 and the temperature to 500°C. The MRM-parameters were optimized using commercially available standards for the SCFAs. The chromatographic separation was performed on a 100 × 2.1 mm, 100 Å, 1.7 μm, Kinetex C18 column (Phenomenex, Aschaffenburg, Germany) column with 0.1% formic acid (eluent A) and 0.1% formic acid in acetonitrile (eluent B) as elution solvents. An injection volume of 1 μL and a flow rate of 0.4 mL/min was used. The gradient elution started at 23% B which was held for 3 min, afterward the concentration was increased to 30% B at 4 min, with another increase to 40%B at 6.5 min, at 7 min 100% B was used which was held for 1 min, at 8.5 min the column was equilibrated at starting conditions. The column oven was set to 40°C and the autosampler to 15°C. Data acquisition and instrumental control were performed with Analyst 1.7 software (Sciex, Darmstadt, Germany).

### Dynamic metabolic profiling of bacterial supernatants

All chemicals were purchased from Sigma Aldrich at the highest purity available. 50 μl of the supernatants were spiked with 100 nmol sodium pyruvate-^13^C_3_ and 250 nmol norvaline as internal standards, afterwards the samples were dried under a gentle stream of nitrogen. For derivatization 100 μl of a methoxyamine hydrochloride solution (10mg/1 ml pyridine) were added and the sample was shaken at 40°C for 90 min. Afterwards 100 μl of MTBSTFA (N-(tert-butyldimethyl-silyl)-N-methyl-trifluoroacetamide containing 1% tert-butyl-dimethyl-silylchlorid) was added and the sample was heated at 70°C for 45 min. GC-MS-analysis was performed with a QP2010 Plus or Ultra gas chromatograph/mass spectrometer (Shimadzu) equipped with a fused silica capillary column (Equity TM-5; 30 m × 0.25 mm, 0.25μm film thickness; SUPELCO) and a quadrupole detector working with electron impact ionization at 70 eV. An aliquot of the derivatized samples was injected in 1:5 split mode at an interface temperature of 260°C and a helium inlet pressure of 70 kPa. After sample injection, the column was first kept at 60°C for 3 min and then developed with a temperature gradient of 10°C min^-1^ to a final temperature of 300°C. This temperature was held for further 3 min.

Pyruvate results were calculated relative to the pyruvate-^13^C_3_ standard (Rt 12.2 min), whereas all other metabolites were calculated relative to norvaline (Rt 17.7 min).

For qualitative sugar analysis 50 μl of the medium were dried under a gentle stream of nitrogen. For derivatization 100 μl of a methoxyamine hydrochloride solution (10mg/1 ml pyridine) were added and the sample was shaken at 40°C for 90 min. Afterwards 100 μl of MSTFA (N-methyl-N (trimethylsilyl)trifluoroacetamide containing 1% trimethylchlorosilane) was added and the sample was heated at 50°C for 45 min. GC-MS-analysis was performed as described above. Glucose, fructose, galactose, mannose and trehalose were confirmed with standard solutions.

### Spot assays

Bacterial cultures and subcultures were grown for 24 hours each in 10ml AF medium at 37°C under anaerobic conditions without shaking. Monocultures were diluted to OD_600nm_ 0.1 in fresh AF medium. To generate a dense bacterial lawn, monoculture inocula were diluted in LB soft agar to OD_600nm_ 0.01 and poured on a AF medium agar plate. After drying all respective other bacteria were spotted onto the bacterial lawn in duplicates in a volume of 5 μl with OD_600nm_ 0.1. Plates were incubated at 37°C for 24h under anaerobic conditions.

### Co-culture experiments

Monoculture inocula were prepared from a previous culture and subculture and were diluted to OD_600nm_ 0.1 in fresh AF medium. Following, pairwise co-cultures were generated by pooling diluted inocula in a 1:1 ratio. From each co-culture 150μl were set aside for pH measurements and determination of initial relative abundances (timepoint 0h). The remaining co-cultures were diluted 1:10 to OD_600nm_ 0.01 and pipetted into a round bottom 96-well plate (Nunc). Growth measurements were performed as described above for 72 hours. Samples for qPCR analysis and pH measurements were taken every 24 hours and the co-cultures were serially diluted 1:100 into 150μl fresh AF medium in a new 96-well round bottom plate to allow communities to approach a steady state composition over ~25 bacterial generations.

### DNA extraction

DNA extraction was performed in the 96-well format using the PureLink^TM^ Pro 96 genomic DNA Kit (Invitrogen) following the corresponding lysis protocol for Gram positive bacterial cells using lysozyme and proteinase K.

### Quantitative PCR of bacterial 16S rRNA genes

Quantitative PCR was performed as described previously (23). Strain-specific 16S rRNA primers and hydrolysis probes were used for amplification. Standard curves were determined using linearized plasmids containing the 16S rRNA gene sequence of the individual strains. The standard specific efficiency was then used for absolute quantification of 16S rRNA gene copy numbers of individual strains.

### Determination of co-culture outcomes

Quantitative 16S rRNA copy numbers from the measurement endpoint of three independent co-culture experiments were determined by qPCR. Co-culture outcomes (positive, neutral or negative) were determined by calculating the individual abundance ratio for each strain in co-culture relative to monoculture. Therefore, the strain specific absolute abundance at 72h in all pairwise co-cultures was divided by the strain specific absolute abundance at 72h in monoculture 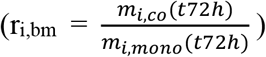 for every individual experiment. Following, the mean abundance ratio from all individual experiments (n=3 per strain combination) was calculated. Significance was determined using a two-sided t-test.

### Community experiments

Monoculture inocula were prepared from a previous culture and subculture and were diluted to OD_600nm_ 0.1 in fresh AF medium. Following, the community inoculum with equivalent ratios of all 12 strains was generated from this dilution. The inoculum was distributed to ten cell culture flasks, thereby diluting the inoculum 1:10 to 10ml fresh AF medium, resulting in a starting OD_600nm_ 0.01. Cell culture flasks were incubated at 37°C without shaking under anaerobic conditions. Every 24h for 10 days samples were taken for qPCR analysis, OD measurement and pH measurement and cultures were diluted 1:100 in 10ml fresh-AF medium.

### Malate uptake measurements

Uptake of ^14^C-malate by *E. faecalis* KB1 was determined in principle as previously described (56). Briefly, *E. faecalis* was grown anaerobically in LB medium with 40 mM malate and harvested in mid-log phase. Cells were centrifuged, washed twice with 50 mM Tris-HCl buffer (pH 7.4) containing 10 mM MgCl_2_ and resuspended in 50 mM Tris-maleate buffer (pH 7.2) containing 5 mM MgCl_2_, thereby adjusting the cell suspension to an OD_600_ of 10. For transport assays, this cell suspension was diluted 1:10 with 50 mM Tris-maleate buffer (pH 7.2) containing 5 mM MgCl_2_ and 1 % (w/v) peptone. Uptake of ^14^C-malate (55 mCi.mmol^-1^ [Biotrend]) was measured at a total substrate concentration of 10 μM at 18° C. At various time intervals, transport was terminated by the addition of stop buffer (100 mM potassium phosphate buffer, pH 6.0, 100 mM LiCl), followed by rapid filtration through membrane filters (MN gf-5 0.4 μm; Macherey-Nagel). The filters were dissolved in 5 ml of scintillation fluid (MP Biomedicals), and radioactivity was determined in a liquid scintillation analyzer (PerkinElmer). Total protein content of *E. faecalis* cells in relation to OD_600_ was determined with a suspension of lysed cells as described before (57).

### Data analysis and Figures

Data was analyzed using R Studio (Version 1.4.1103). Heatmaps were generated using the R *pheatmap* package (https://github.com/raivokolde/pheatmap). Plots were generated using the R *ggplot2* package (58) and *ggpubR* package (https://github.com/kassambara/ggpubr). Figures were partly generated using BioRender (https://biorender.com) and Adobe Illustrator CC (Adobe Inc.).

## Acknowledgements

The authors thank D. Ring, C. Beck and C. Schwarz for their technical support and members of the Stecher laboratory for helpful feedback and discussions. This research received funding by the German Research Foundation (DFG, German Research Foundation, Projektnummer 395357507–SFB 1371, Projektnummer 279971426 and 315980449), the European Research Council (ERC) under the European Union’s Horizon 2020 research and innovation programm (Grant Agreement 865615), the German Center for Infection Research (DZIF) and the Center for Gastrointestinal Microbiome Research (CEGIMIR).

## Author contributions

B. S., A.S., K.J. and A.S.W. conceived and designed the experiments. A.S.W., A.v.S., A.B., A.C.D.R., L.R., S.G., K.K. and C.H. performed the experiments. A.S.W., A.B., A.C.D.R., L.R., C. M. C.H., K.K., S.G. and P.M. analyzed the data. P.M., K.K., C.M., C.H., K.J. and W.E. contributed materials/ analysis tools. B.S. coordinated the project. B.S., L.M.J. and A.S.W. wrote the original draft and all authors reviewed and edited the draft manuscript.

