## Supplementary Information for "Exploring the interaction network of a synthetic gut bacterial community"

#### Supplemental Figures

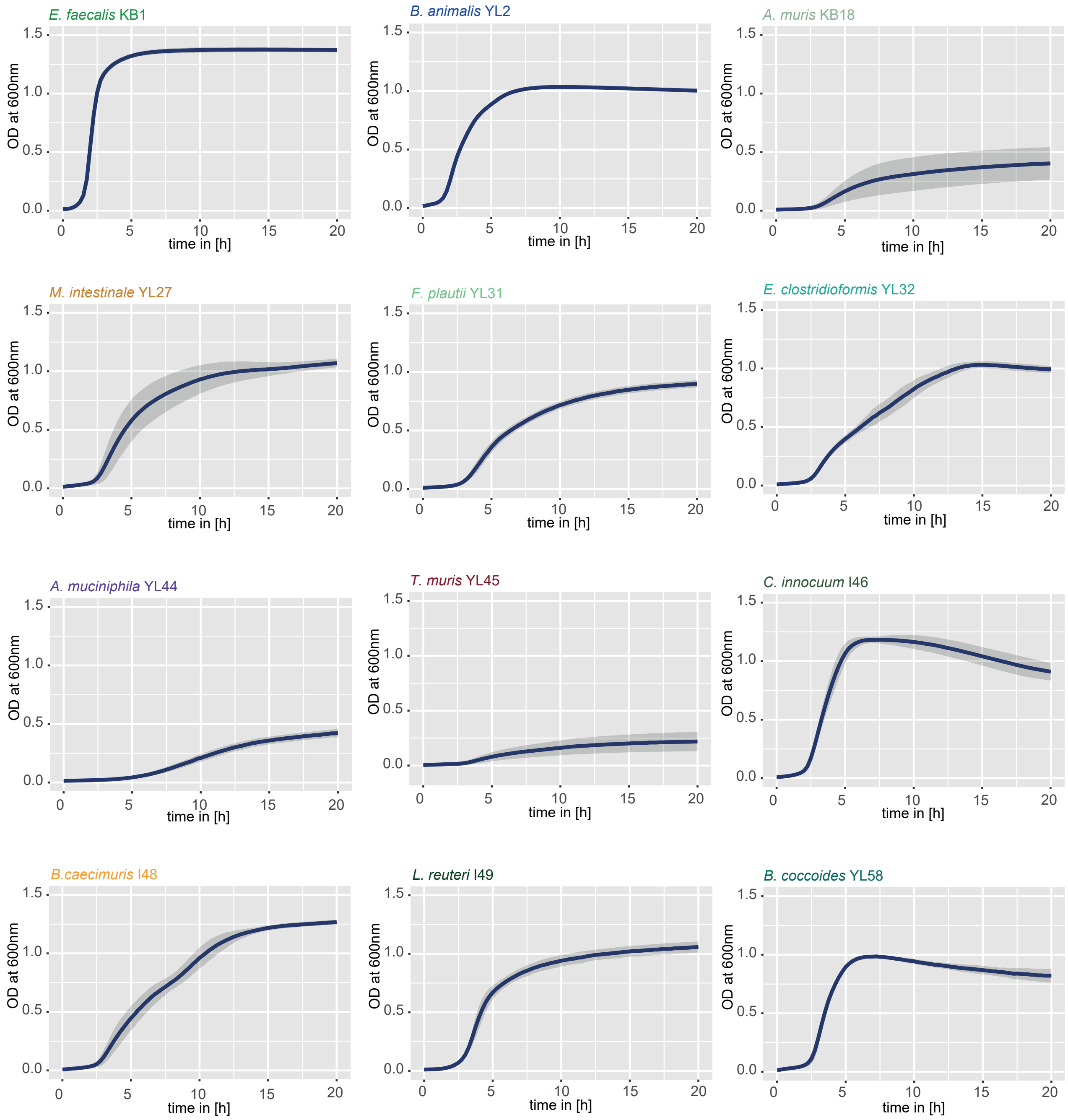


**Fig. S1. Growth characteristics of individual strains in monoculture.** Growth in AF-medium was monitored at OD 600nm, mean (blue line) and SD (grey) of three independent experiments is shown. Growth rates were extrapolated from growth in the exponential phase (Tab. S3). All strains grew to stationary phase within 20 hours and strain specific behavior in AF-medium can be observed. Interestingly, strains *E. clostridioformis* YL32 and *B. caecimuris* I48 show indications of a diauxic growth behavior. Further, strains *C. innocuum* I46 and *B. coccoides* YL58 show a decrease in OD 600nm after stationary phase is reached.

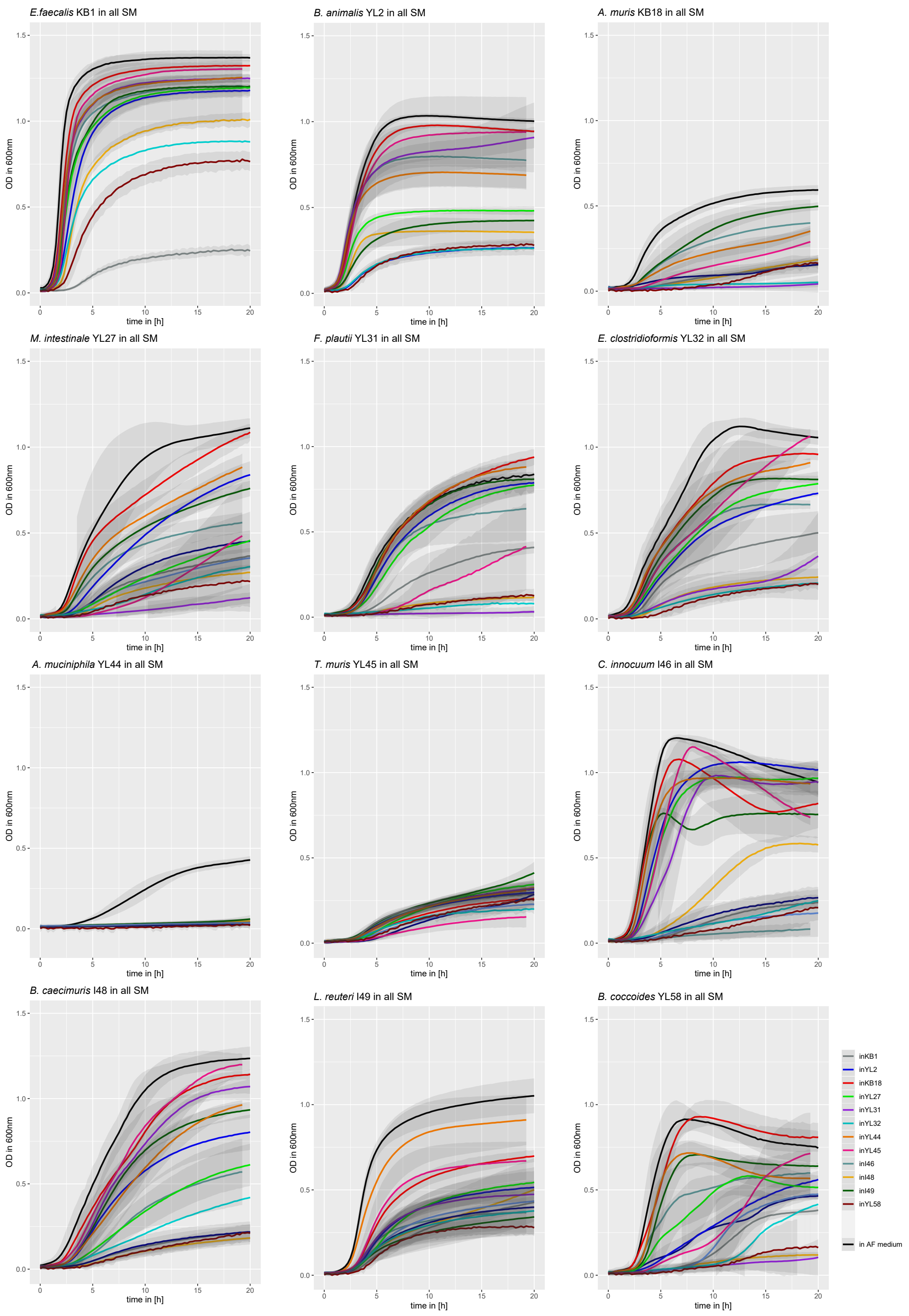


**Fig. S2. Growth curves of individual strains in SM of all OMM[12] bacteria.** Growth of all individual monocultures was monitored in fresh AF-medium (black) and in SM of individual OMM[12] strains (colored lines) over 20 hours at OD 600nm. The mean of three indepedent experiments is shown with the corresponding SD (grey).

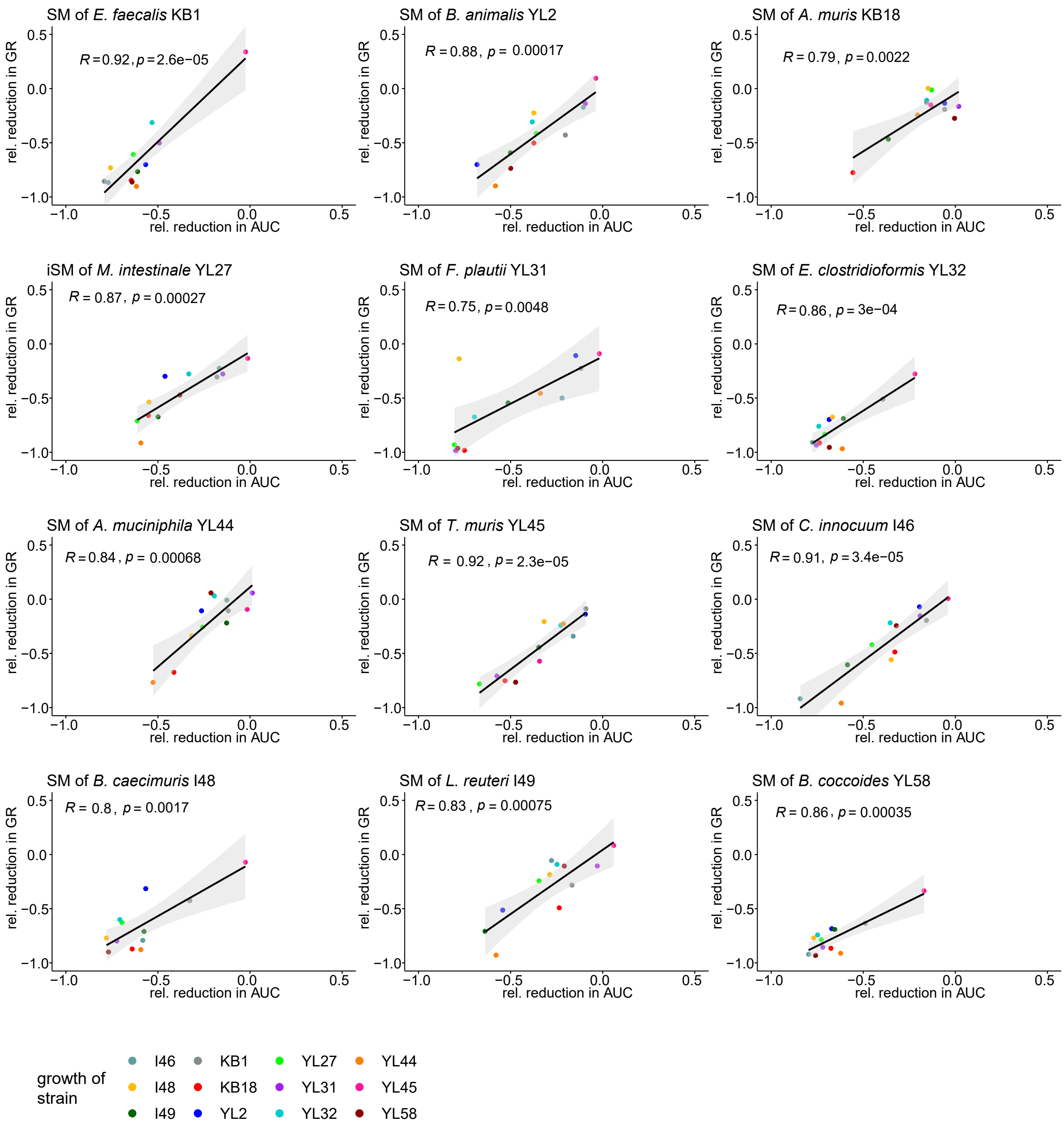


**Fig. S 3. Correlation of reduction in growth rate and reduction in AUC.** The reduction of growth rate and AUC in SM relative to the corresponding fresh AF medium control was calculated for all strains of the OMM[12] consortium from data shown in **Fig S2**. Relative coefficients were then correlated and show a negative linear relation for all individual strains, indicating that both growth rate and AUC in a specific SM are decreased to the same extent. The corresponding values for the individual strains are shown in colors, the linear fit (black) with the standard error (grey).

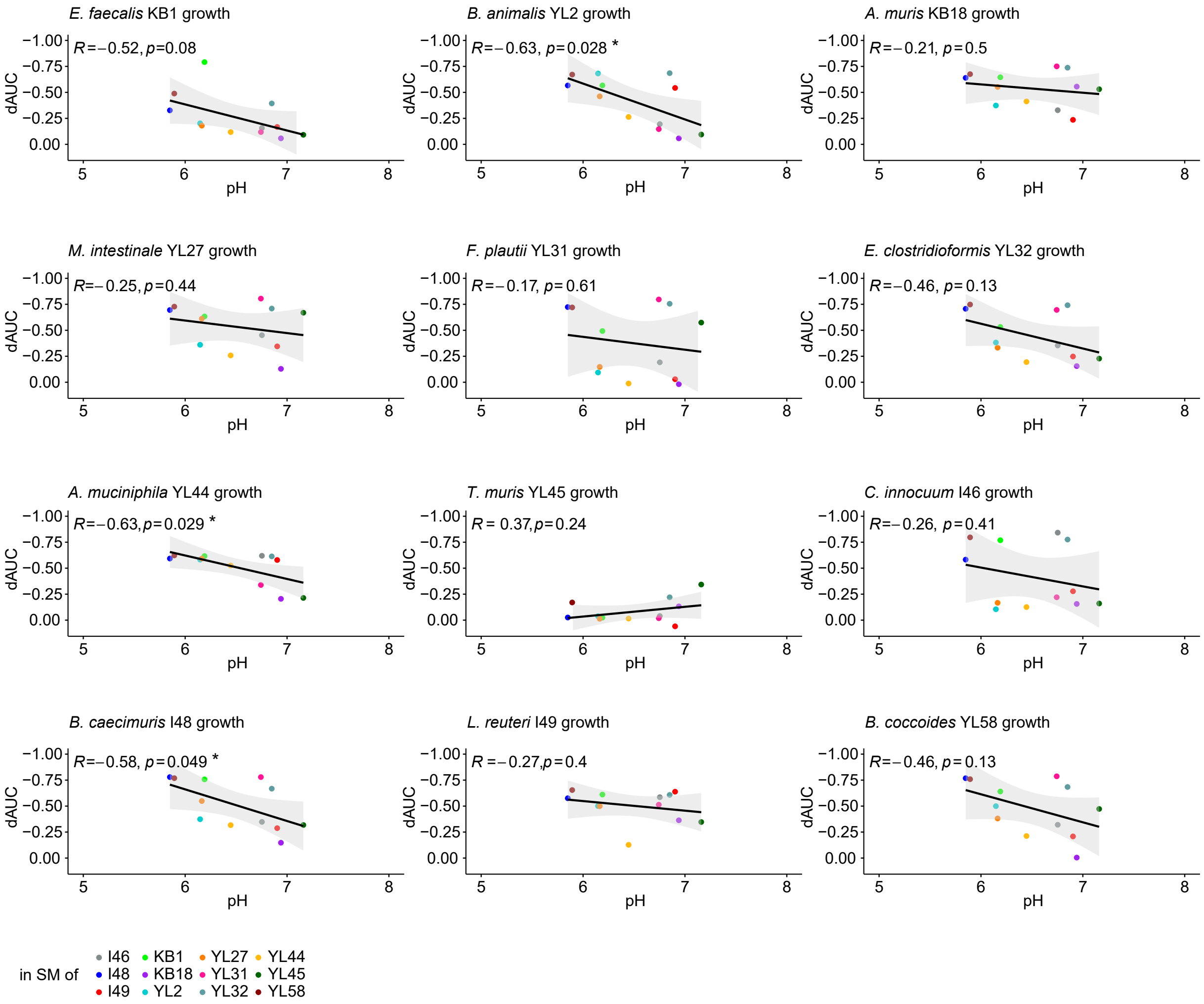


**Fig. S4. Correlation of reduction in AUC with the pH of the SM of a specific strain.** Correlation of the inhibition of growth in a SM ($d_{AUC}$) with the mean pH of the individual SM for each strain revealed that growth inhibition does not directly correlate with the pH for all strains. Only *B. animalis* YL2, *A. muciniphila* YL44 and *B. caecimuris* I48 showed a significant negative correlation ($R < -0.5$, $p < 0.05$) between growth inhibition and pH with stronger inhibition at more acidic pH ranges. The corresponding values for the individual strains are shown in colors, the linear fit (black) is depicted with the SD (grey).

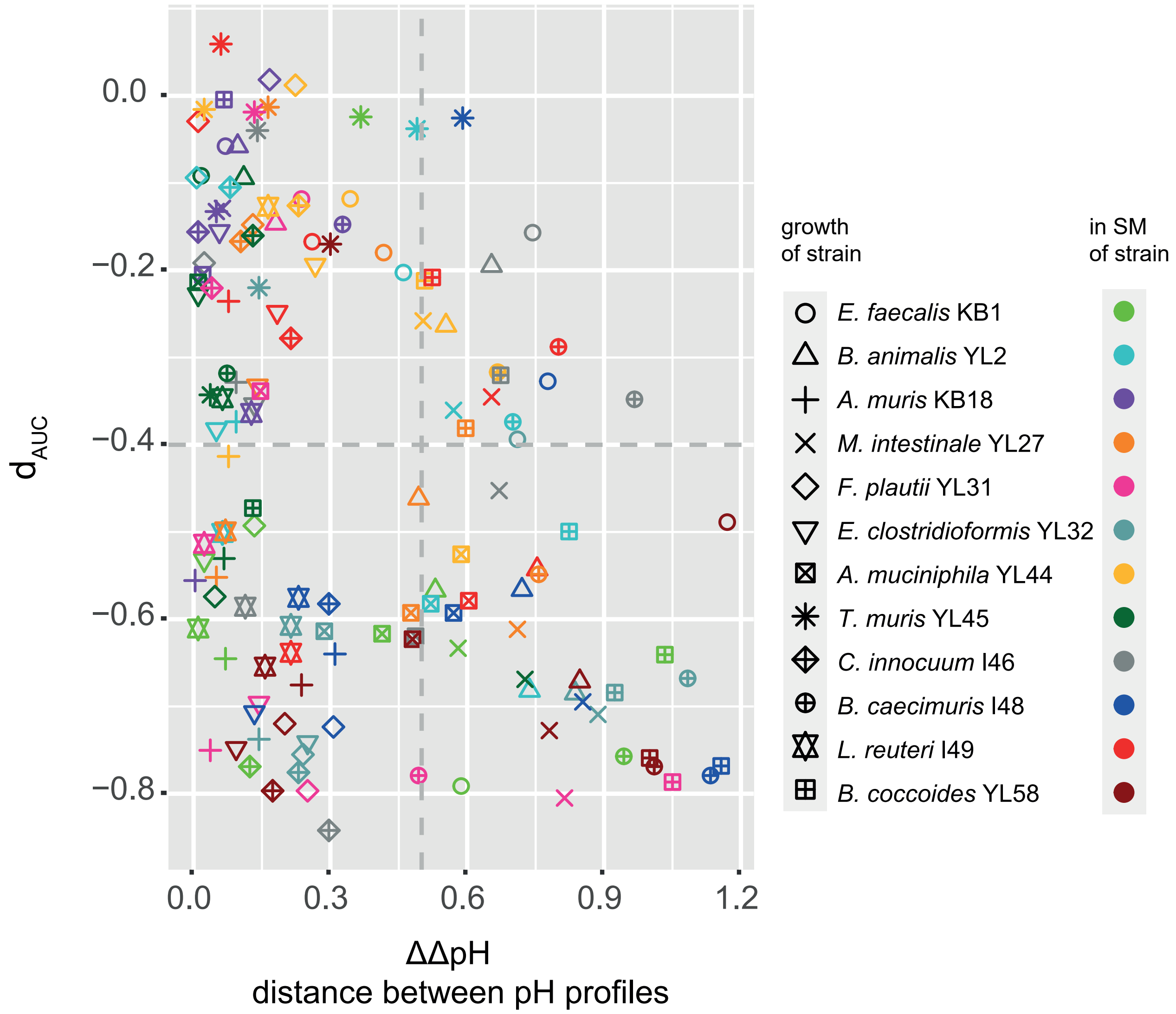


**Fig. S5. Relationship between AUC decrease in SM and change in pH profiles.** The factor ΔΔpH depicts the Euclidean distance between the $\Delta pH_{SM}$ (after growth in fresh AF medium) and the corresponding $\Delta pH_{DSM}$ (after growth in a respective SM) and is a measure how the pH profile of a individual strain changes in the environment of a specific SM in comparison to fresh medium (see supplemental text). Cases where overall growth is only slightly influenced in SM ($d_{AUC} > -0.4$), but the corresponding ΔpH changed strongly ($\Delta\Delta pH > 0.5$, upper right quadrant) might indicate a drastic change in metabolic behavior of a strain due to the altered environmental conditions in SM. For example, the ΔΔpH of *E. faecalis* KB1 in the SM of *C. innocuum* I46 indicates (grey circle), that here the metabolic behavior of KB1 is altered in comparison to fresh AF medium. While KB1 strongly acidifies the neutral fresh culture medium($pH = 7.0$) to $pH_{SM,\ KB1} = 6.19$ ($\Delta pH_{SM} =$ - 0.81), the neutral SM of I46 ($pH_{SM,\ I46} =$ 6.75) is not distinctly acidified after KB1 growth ($pH_{DSM} = 6.68$, corresponding to $\Delta pH_{DSM} =$ - 0.07 and therefore $\Delta\Delta pH = 0.74$).

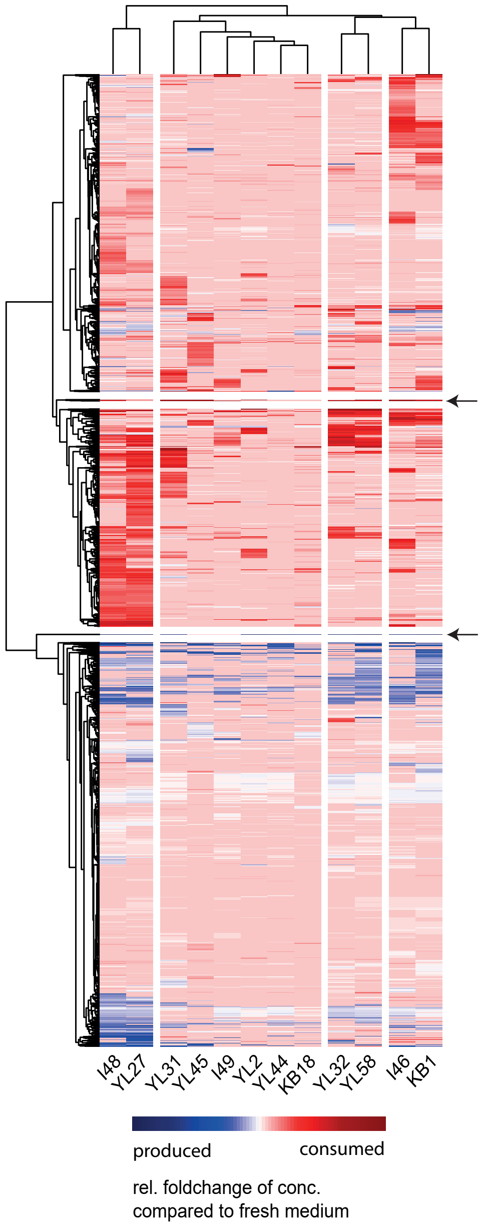


**Fig. S6. Metabolomic analysis of SM.** Metabolomic profiles after bacterial growth to stationary phase in AF medium were determined by untargeted MS (methods). All metabolomic features that significantly changed for at least one of the twelve strains are shown (rows). Levels decreased (red) and levels increased (blue) compared to fresh AF medium as determined by the relative foldchange. Hierarchical clustering of strain specific profiles as well as measured metabolomic features reveal profile similarities between phylogenetically related strains. Black arrows indicate two separated clusters of metabolomic features that are produced and consumed by most bacterial strains.

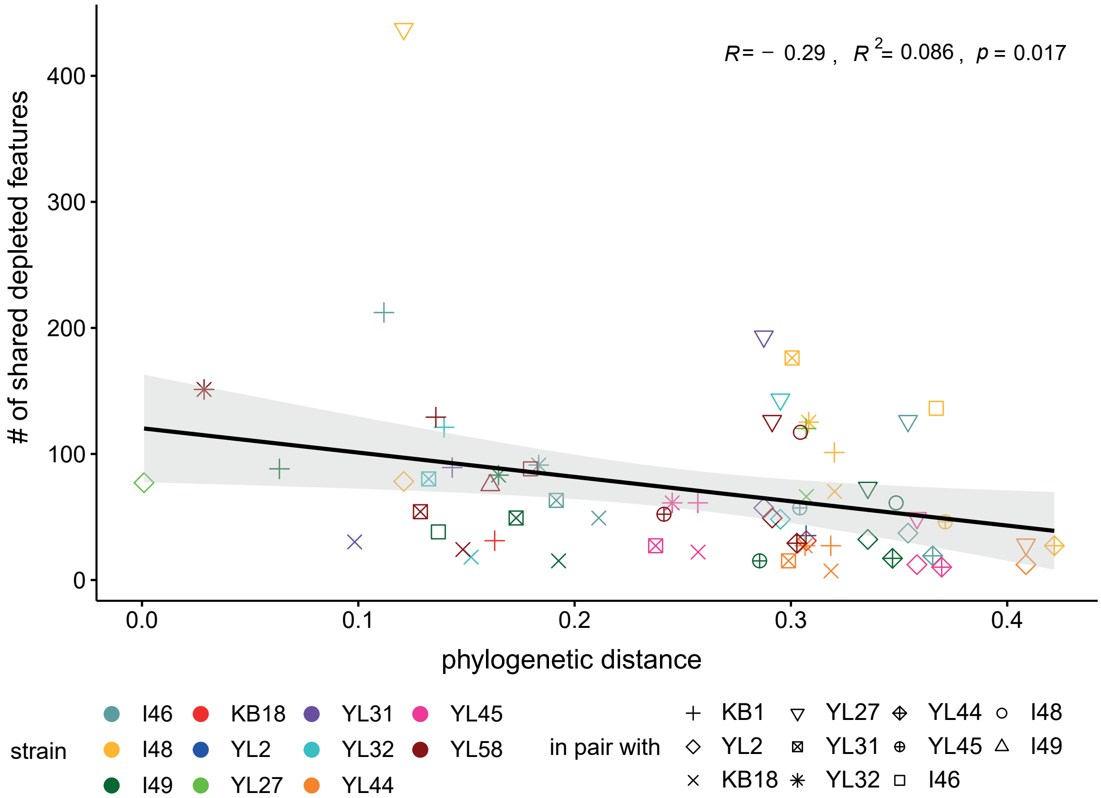


**Fig. S7. Correlation of overlap in substrate depletion profiles with phylogenetic distance.** Correlating the phylogenetic distance between the individual strains (based on 16S rRNA gene sequences) with the number of shared depleted metabolic features in AF medium showed that phylogenetically similar strains of the consortium have a higher substrate overlap than phylogenetically distant strains. The corresponding values for the individual strain pairs are shown in colors and shapes, the linear fit is depicted (black line) with the standard error (grey).

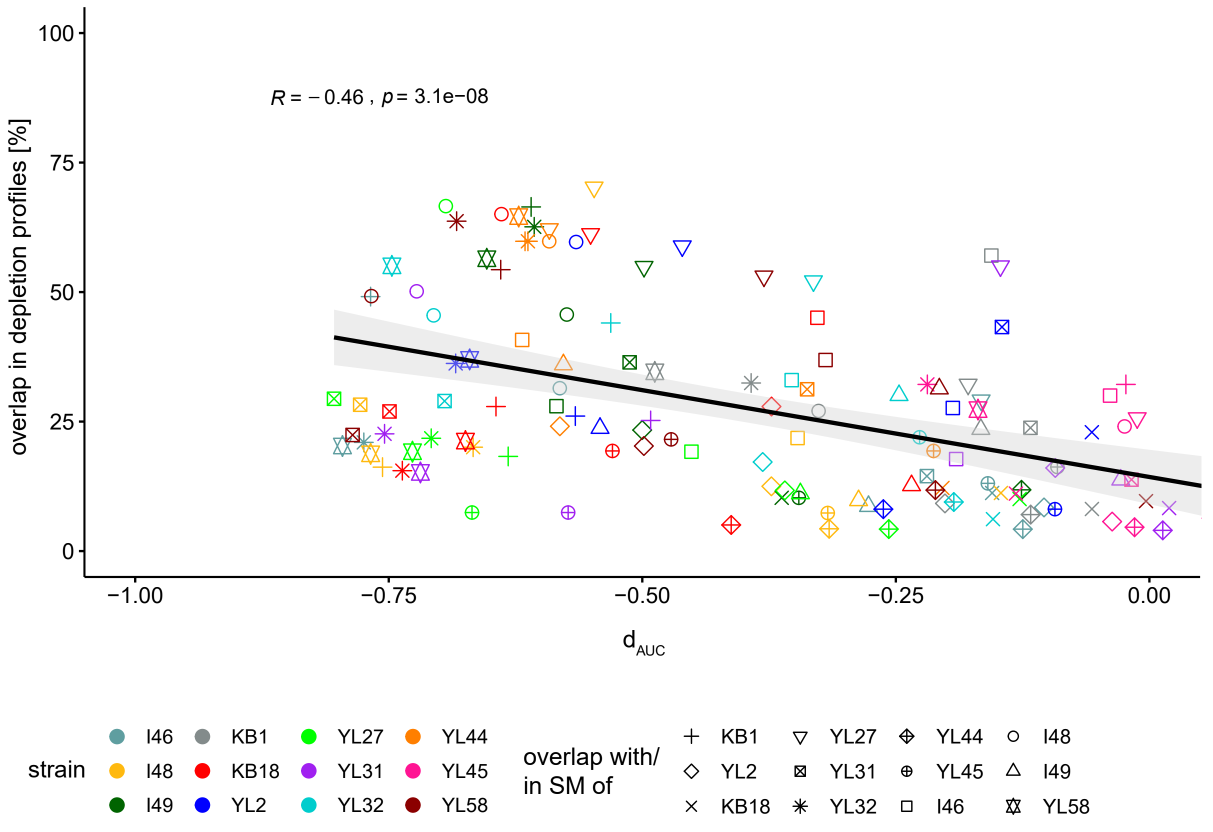


**Fig. S8. Correlation of overlap in substrate depletion profiles with inhibition of growth in SM.** Correlating the pairwise overlap in depleted metabolic features in AF medium with the inhibition of growth in the corresponding SM ($d_{AUC}$) revealed that overlap in depletion profiles is correlated with growth inhibition in the corresponding SM. The corresponding values for the individual strain pairs are shown in colors and shapes, the linear fit is depicted (black line) with the standard error (grey).

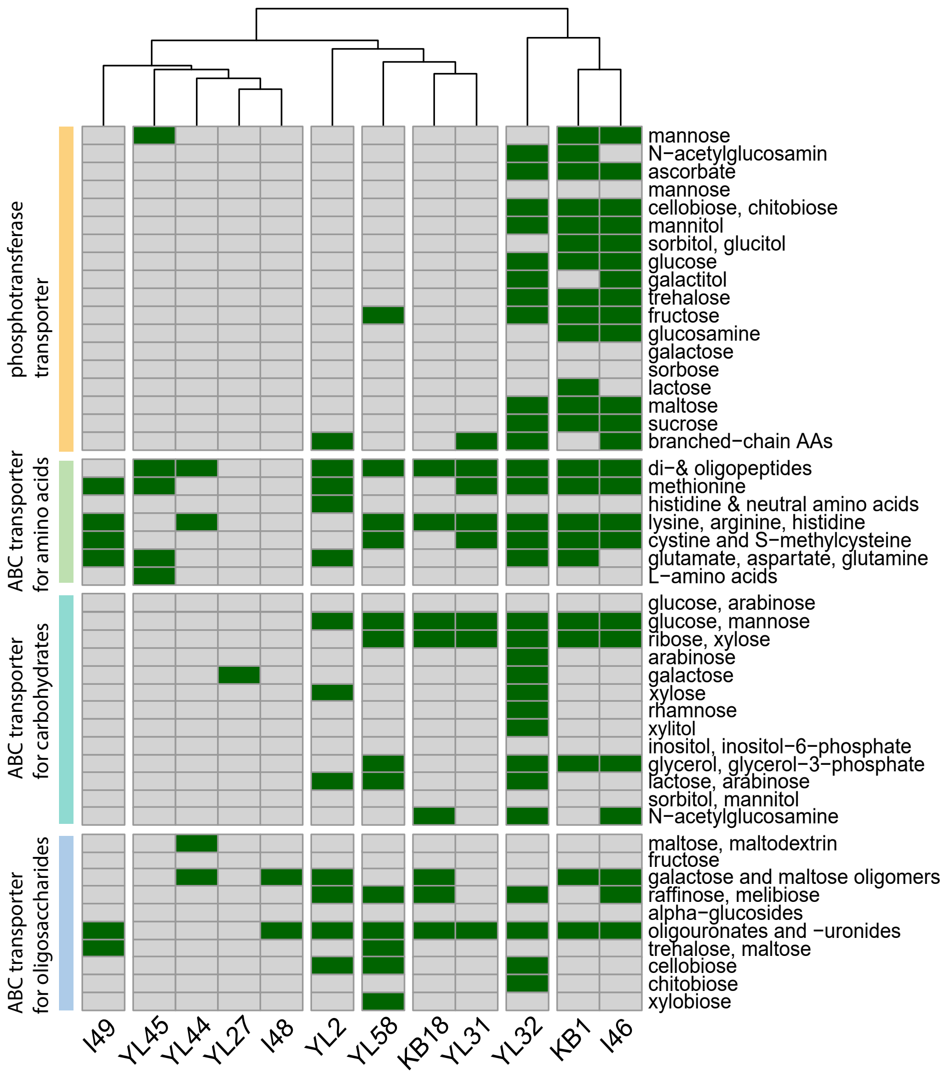


**Fig. S9. Genome-informed potential for substrate transport.** Different groups of substrate specific transporters (ABC transporters and PTS-systems) were specified and genome KO annotations were mined for the corresponding key enzymes (SI data table). A positive hit (green) does not indicate the presence of a complete pathway, but at least one key KO annotation. Hierarchical clustering of strain specific profiles revealed profile similarities especially between strains *E. faecalis* KB1, *C. innocuum* I46 and *E. clostridioformis* YL32.

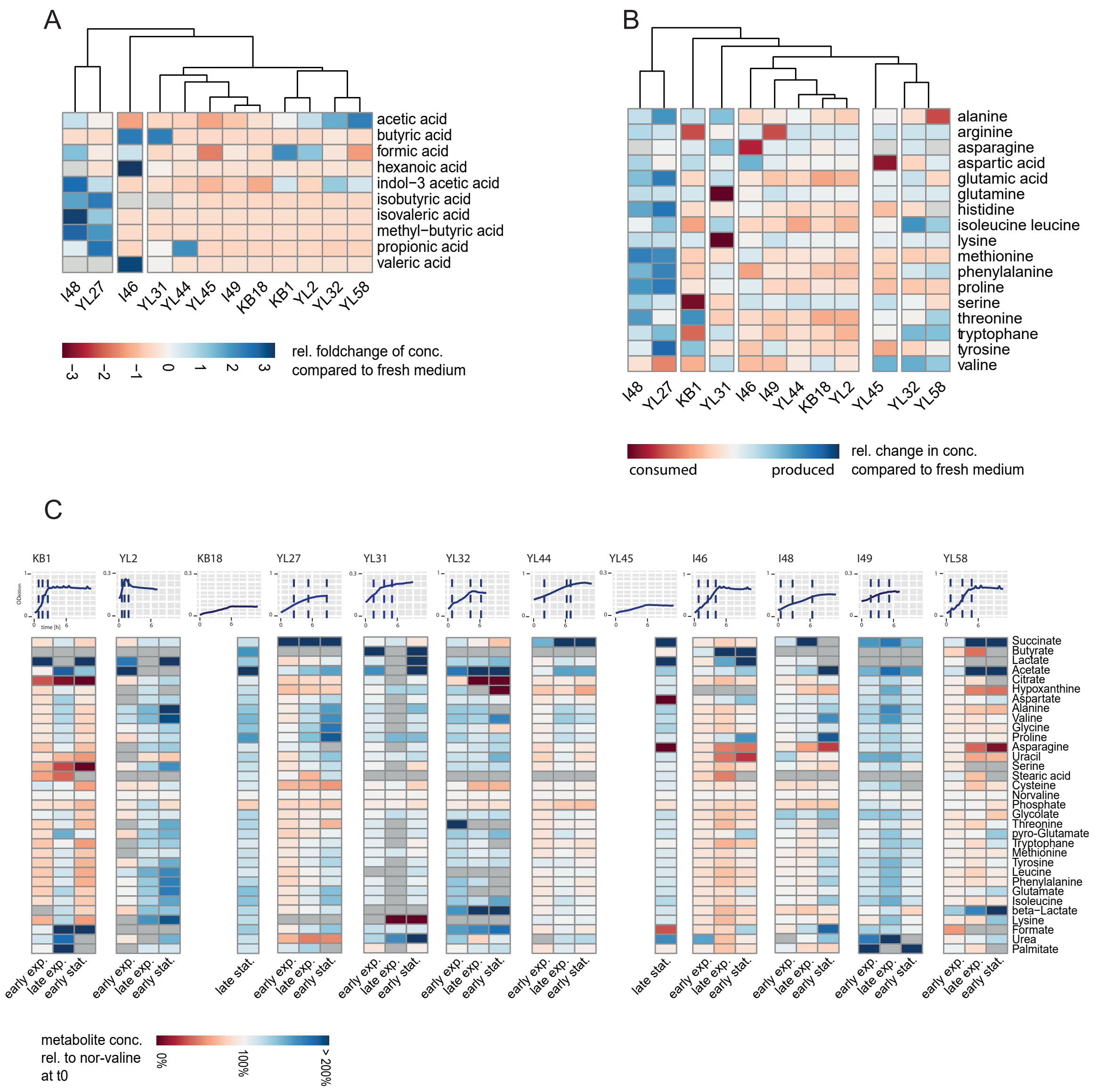


**Fig. S 10. Metabolic profiling of SM.** (**A**) SCFA profiles as determined by targeted metabolomics analysis of SM (late stationary phase, methods). SCFA consumption is indicated in red, while production is indicated in blue. (**B**) Amino acid profiles as determined by untargeted metabolomics analysis of SM (late stationary phase). Amino acids were annotated by comparing the exact mass and MS2 fragmentation patterns of the measured features to the records in HMBD (methods). Consumption of a specific amino acid from fresh medium is indicated in red, while production in indicated in blue. (**C**) Strain-specific metabolite profiles in different growth phases (early exponential, late exponential and early stationary phase) by GC-MS. Bacterial supernatants were sampled while simultaneously monitoring the individual growth stages by measuring OD at 600nm. Using an internal norvaline standard it was determined which metabolite levels increase (blue) and decrease (red) over time.

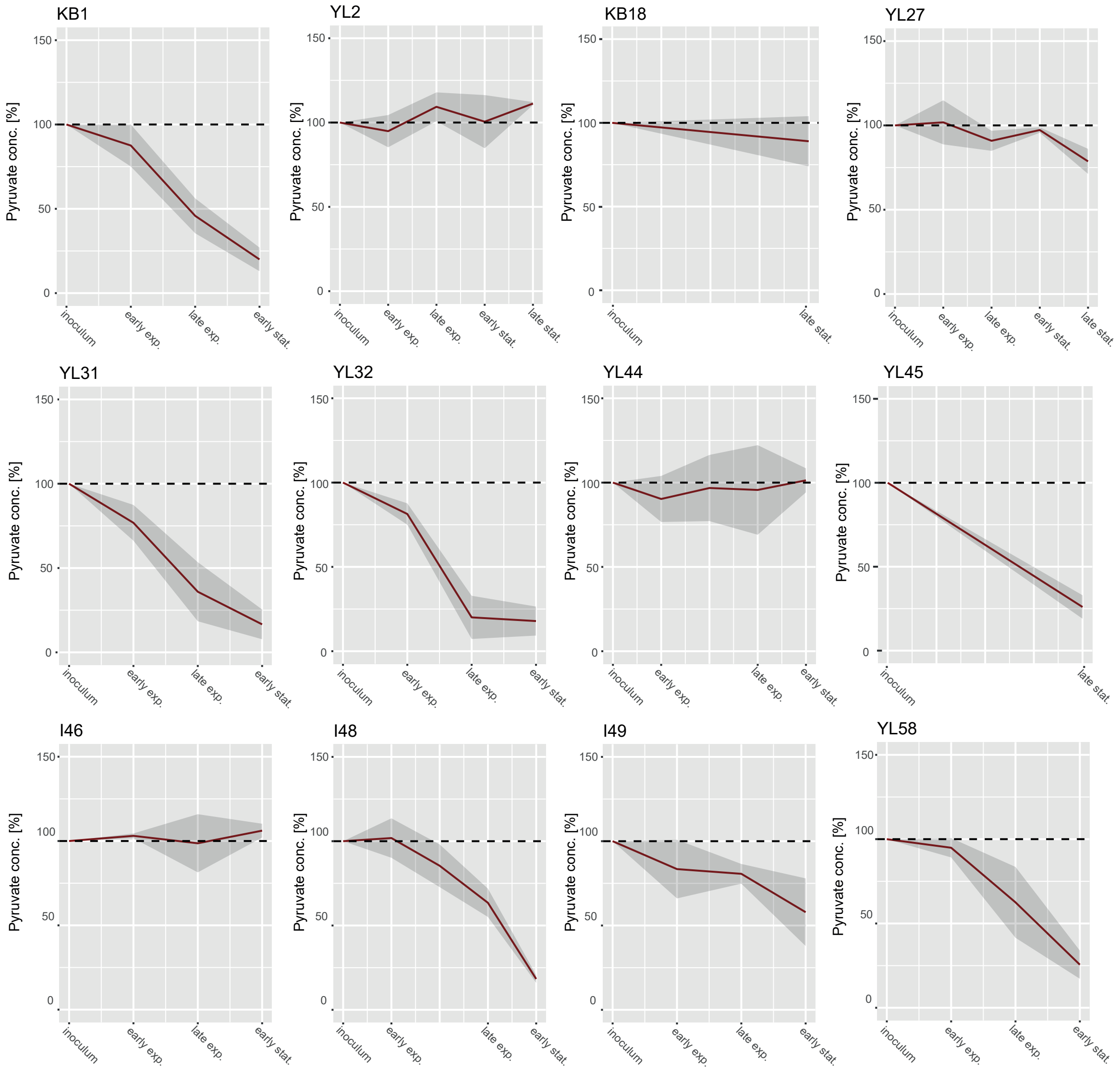


**Fig. S 11. Dynamic probing of pyruvate levels.** Bacterial supernatants sampled at different time points of monoculture growth were spiked with $^{13}C$ labeled sodium pyruvate by GC-MS. Mean pyruvate levels (red line) are shown relative to fresh AF-medium in percent with the corresponding standard deviation shown in grey.

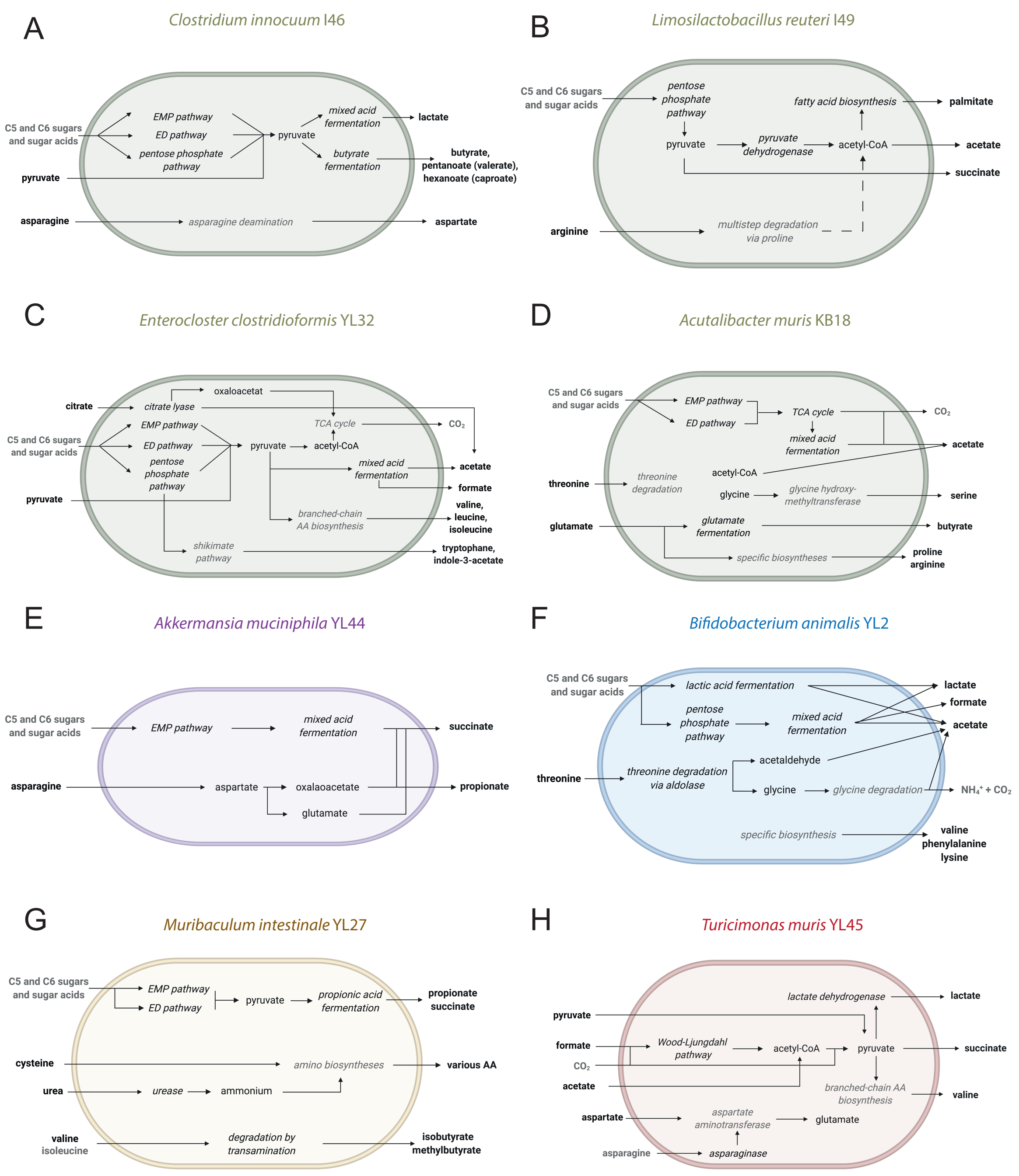


**Fig. S12. Draft metabolic models of the individual OMM$^{12}$ community members.** Combining metabolomics analyses with genome-based information on the presence of key enzymes enabled the generation of broad-scale draft metabolic models of the individual OMM$^{12}$ community members. Experimentally confirmed substrates, products, enzymes or pathways are shown in black. Hypothetical substrates, products, enzymes or pathways are shown in grey. Substrates, intermediate products or products are shown in roman, while enzymes or pathways are shown in italic font. Extracellular substrates or products are shown in bold. Additional information on the individual broad scale draft metabolic models can be found in the supplementary text.

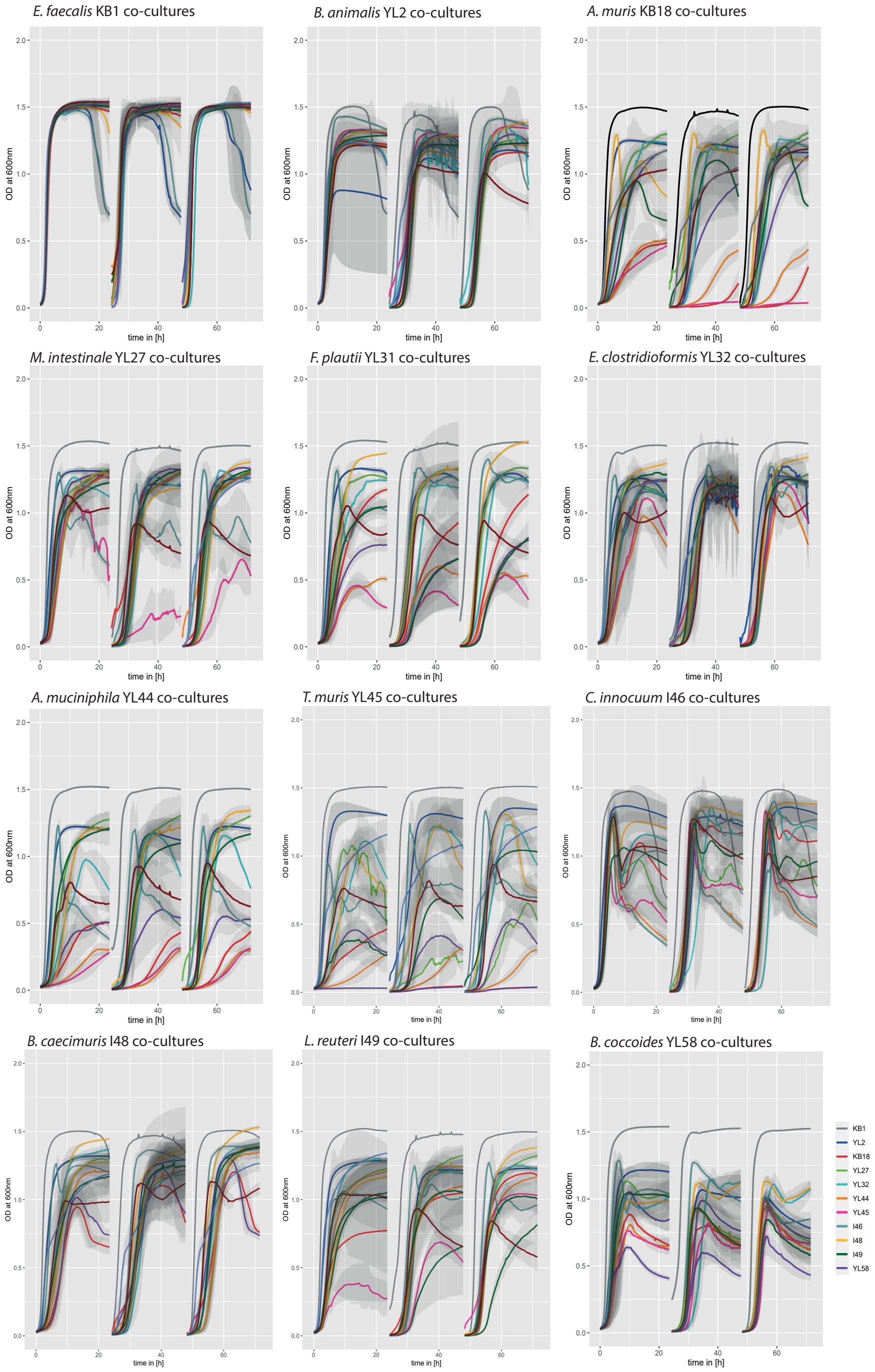


**Fig. S13. Growth curves of all OMM$^{12}$ bacteria in co-culture with all respective other strains.** Growth of all co-cultures was monitored over 20 hours at OD 600nm. The corresponding monoculture is shown in black, while all respective co-cultures are shown in colored lines and standard deviations shown in light grey.

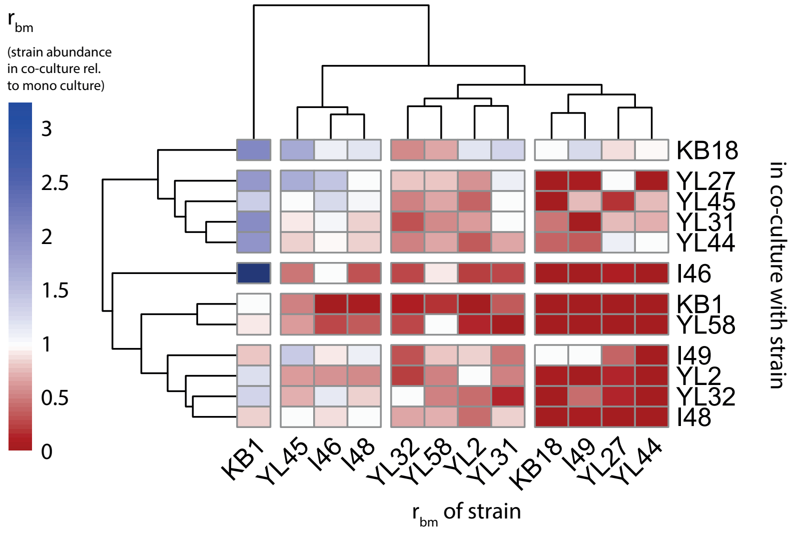


**Fig. S14. Strain specific absolute abundance in pairwise co-culture relative to monoculture.** Mean absolute abundance (16S rRNA gene copies determined by qPCR) ratio $r_{bm}$ as a measure of how successful a strain can grow in co-culture relative to monoculture after 72h. A ratio $r_{bm} = 1$ indicates no change in absolute abundance in the co-culture compared to mono culture. A ratio $r_{bm} > 1$ and a ratio $r_{bm} < 1$ indicate an increase and decrease in absolute abundance in the co-culture compared to mono culture, respectively.

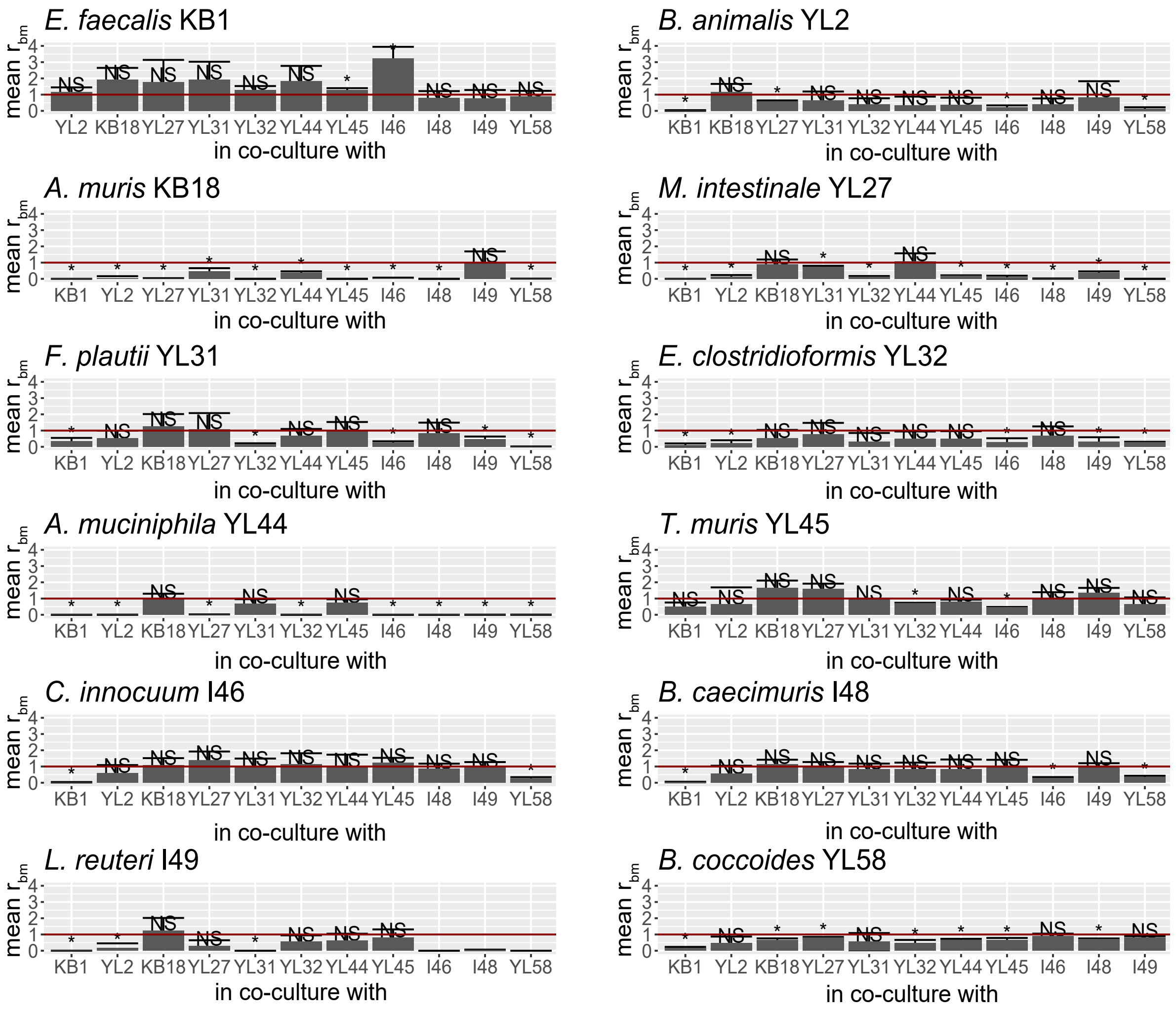


**Fig. S15. Significance analysis of mean $r_{bm}$ in all co-cultures.** Mean absolute abundance ratios $r_{bm}$ are shown as barplot with the corresponding standard deviation. The reference value (equal to one) is shown as red horizontal lines. A t-test was performed to determine significant increase or decrease in $r_{bm}$ in the individual co-cultures relative to monoculture ($r_{bm}$ =1). Significantly changed absolute abundance ratios $r_{bm}$ ($p < 0.05$) are indicated by (*), non-significantly changed absolute abundance ratios $r_{bm}$ are indicated by (NS).

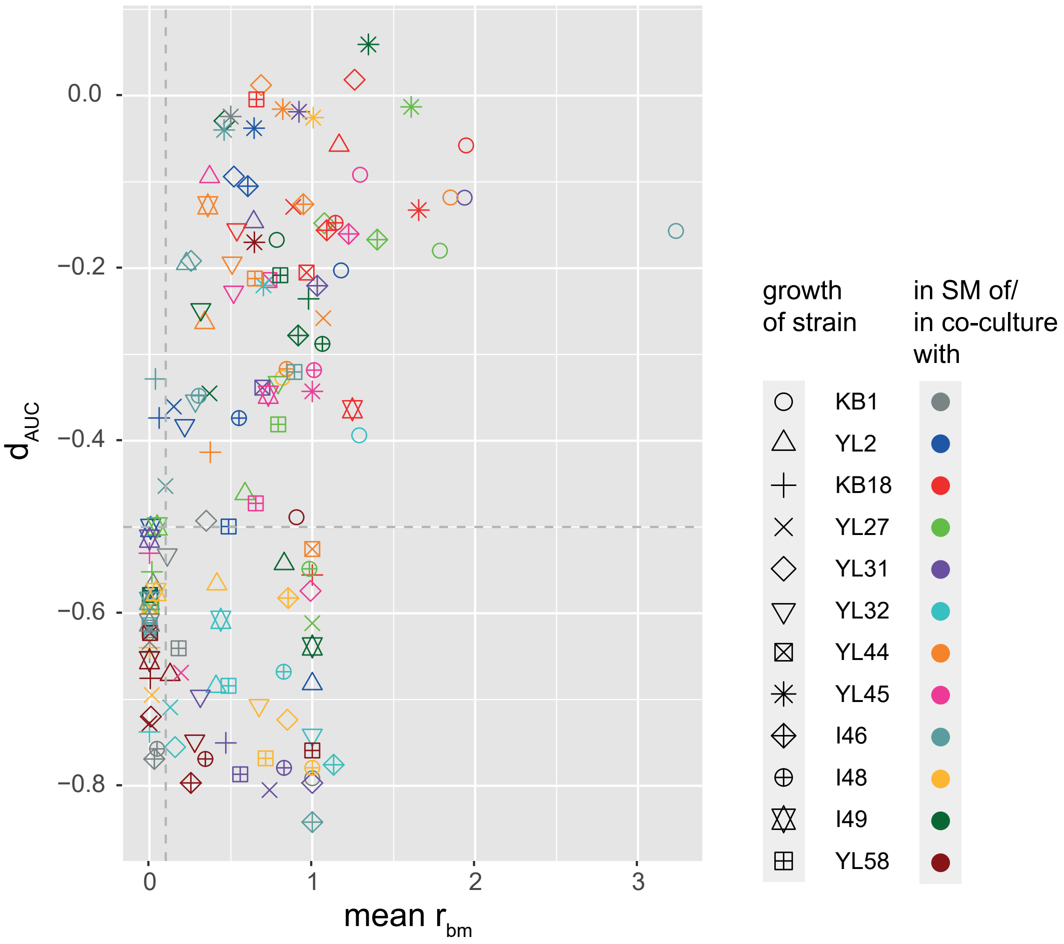


**Fig. S16. Relationship between strain specific influence by SM and in co-culture.** Comparing the influence of a strain on others in co-culture growth ($r_{bm}$) with the degree to which other strains were inhibited by its SM ($d_{AUC}$). Most co-cultures that resulted in the extinction of strain i, while strain j was not or positively affected ($r_{i,bm} \approx 0$, while $r_{j,bm} \geq 1$ ) also showed high inhibition values of strain i in the SM of strain j ($d_{i,AUC} \leq$ -0.5). This suggests, that in most cases a strongly negative co-culture outcome for strain i corresponds to a strong inhibition of strain i by strain j due to specific waste or end products or substrate overlap.

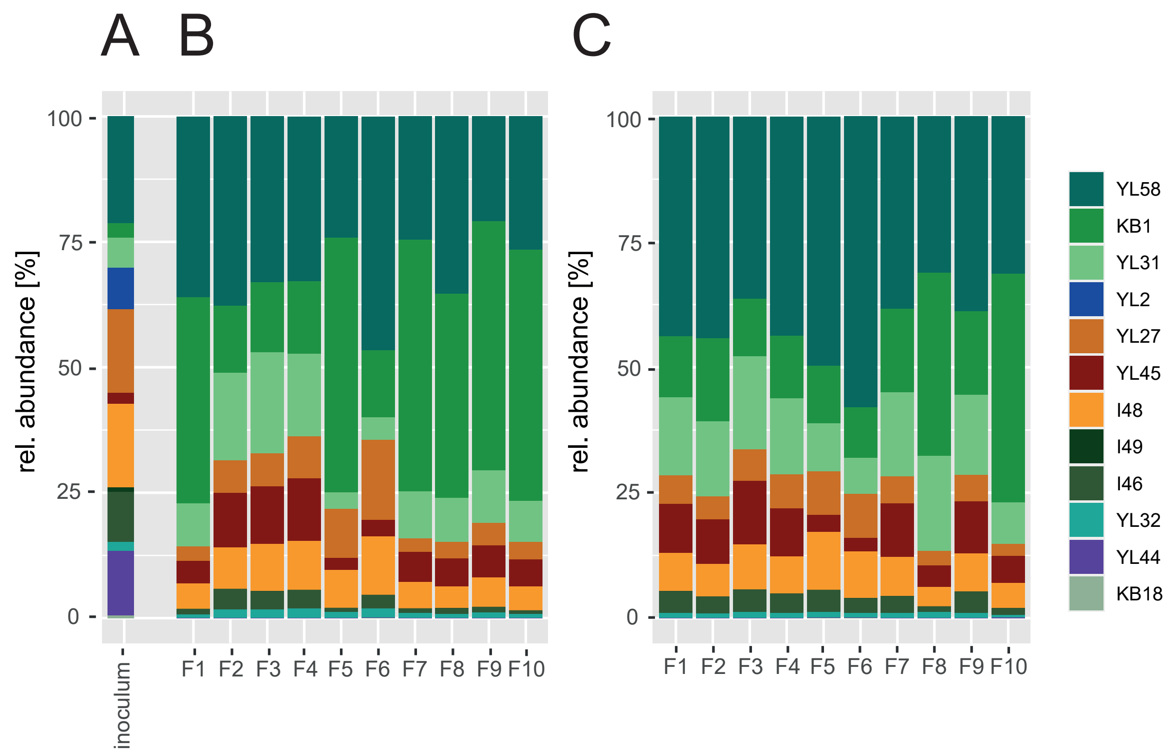


**Fig. S17. Relative abundance of the OMM$^{12}$ strains in community batch culture.** Relative abundance of all 12 strains as determined by qPCR is shown for the inoculum (**A**), all replicates (F1-F10) after three days (**B**) and ten days (**C**) of semi-continuous culture. While strains *B. animalis* YL2 and *L. reuteri* I49 were below detection limit in all replicates, strain *A. muris* KB18 was detectable in two of the ten replicates (relative abundance $< 1\%$). Data shown here was generated in an independent replication of the experiment shown in Fig. 5. Even though the inoculum differs slightly in its composition, the community structure approaches the same composition in both experiments.

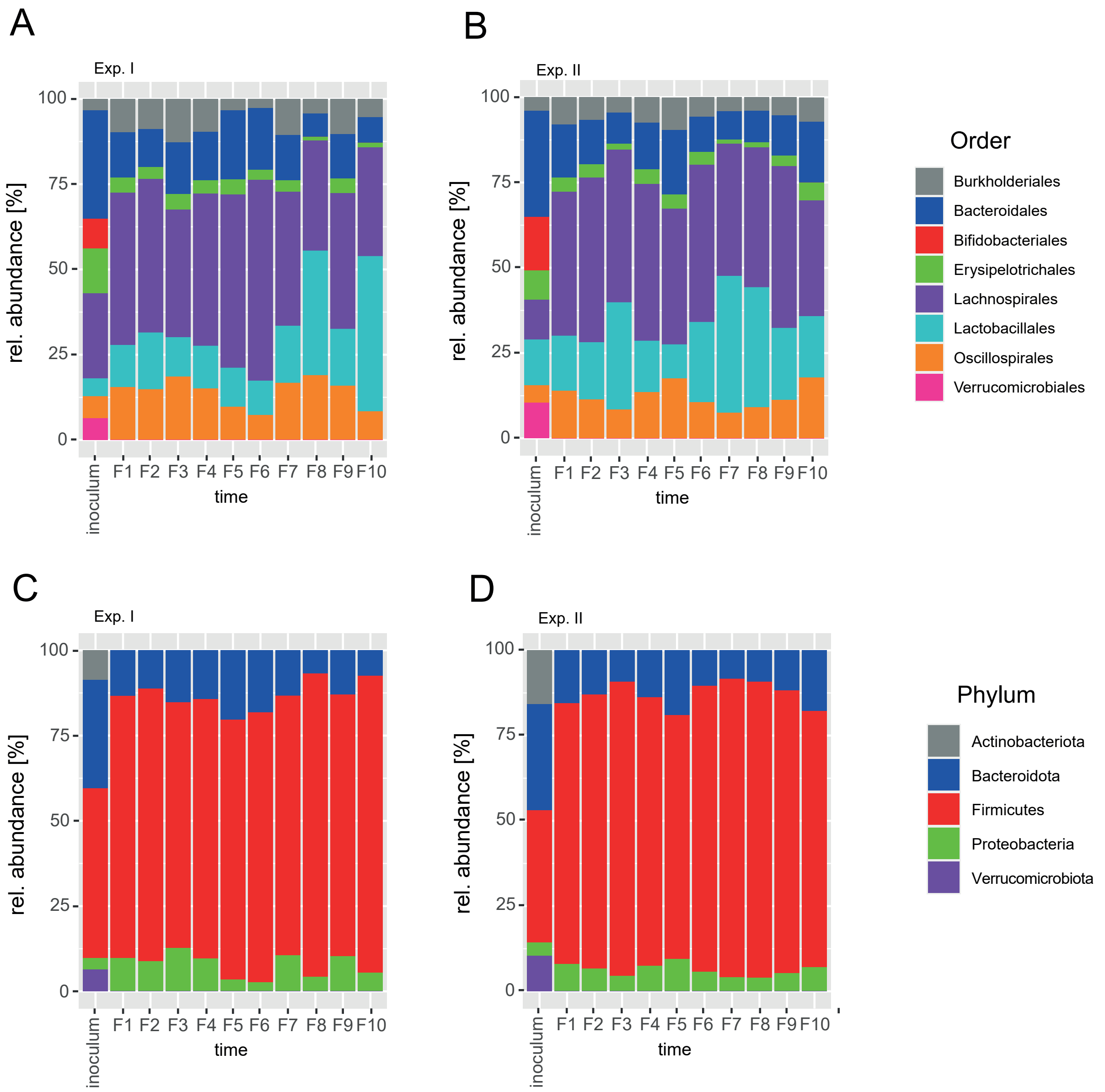


**Fig. S18. Relative abundance of the OMM$^{12}$ strains in community batch culture on the order and phylum level.** Relative abundance of all 12 strains as determined by qPCR is shown for the inoculum and all replicates (F1-F10) after ten days of semi-continuous batch culture. Data was generated in two independent experiments (I II) with cultures from the same inoculum. Relative abundance on the respective order and phylum level of experiment I in (**A**) and (**C**) and of experiment II in(**B**) and (**D**).

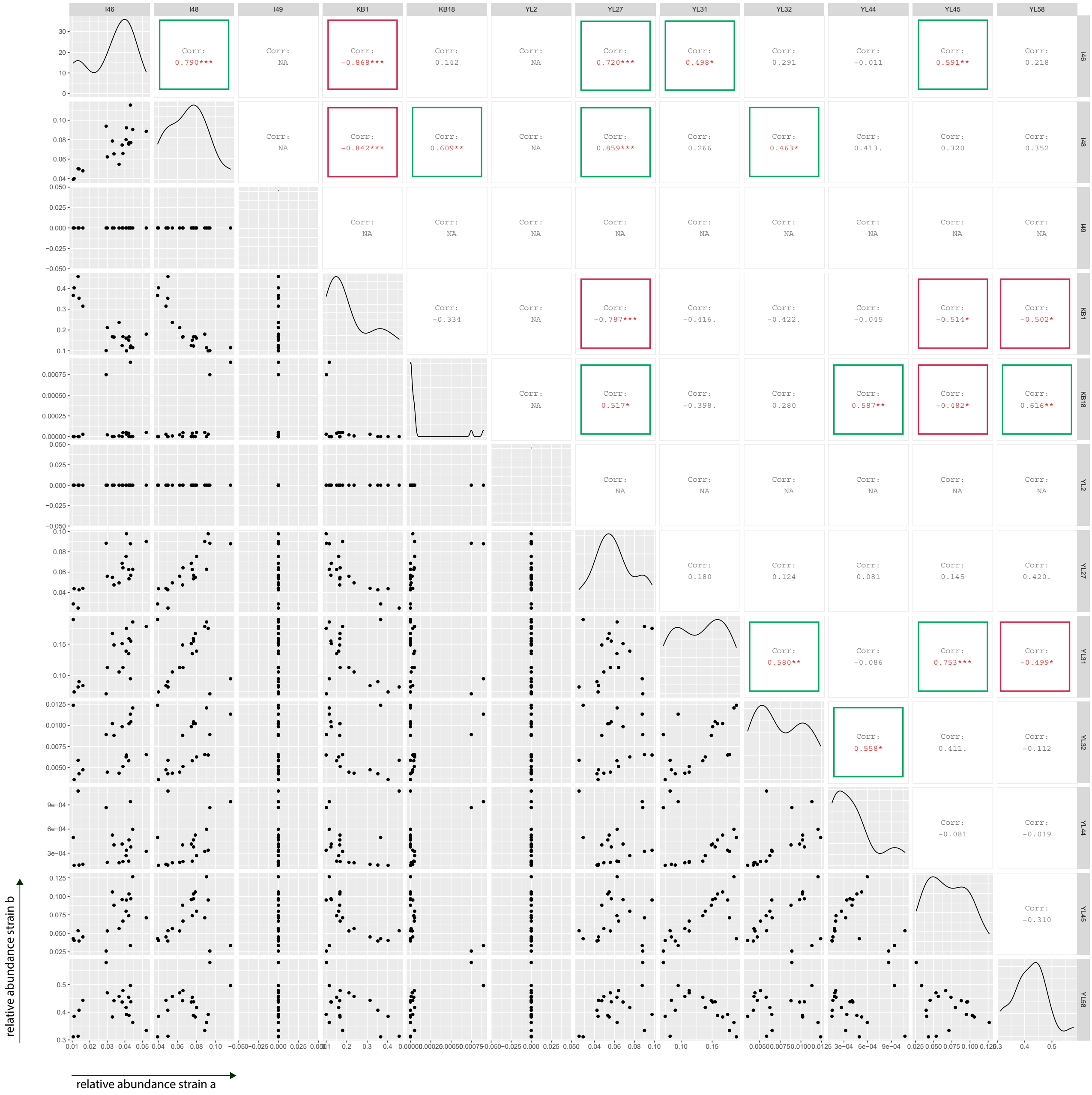


**Fig. S19. Pairwise correlation of strain specific relationships in community batch culture from relative abundance.** Using the information on relative abundance of the twelve strains in community batch culture from two experiments with ten replicates each, pairwise strain relationships were determined, performing a pearson correlation (*ggcorr* in the *GGally* package in R). Significant correlation coefficients are highlighted in red text. Positive correlations are marked with a green, negative correlations with a red frame.

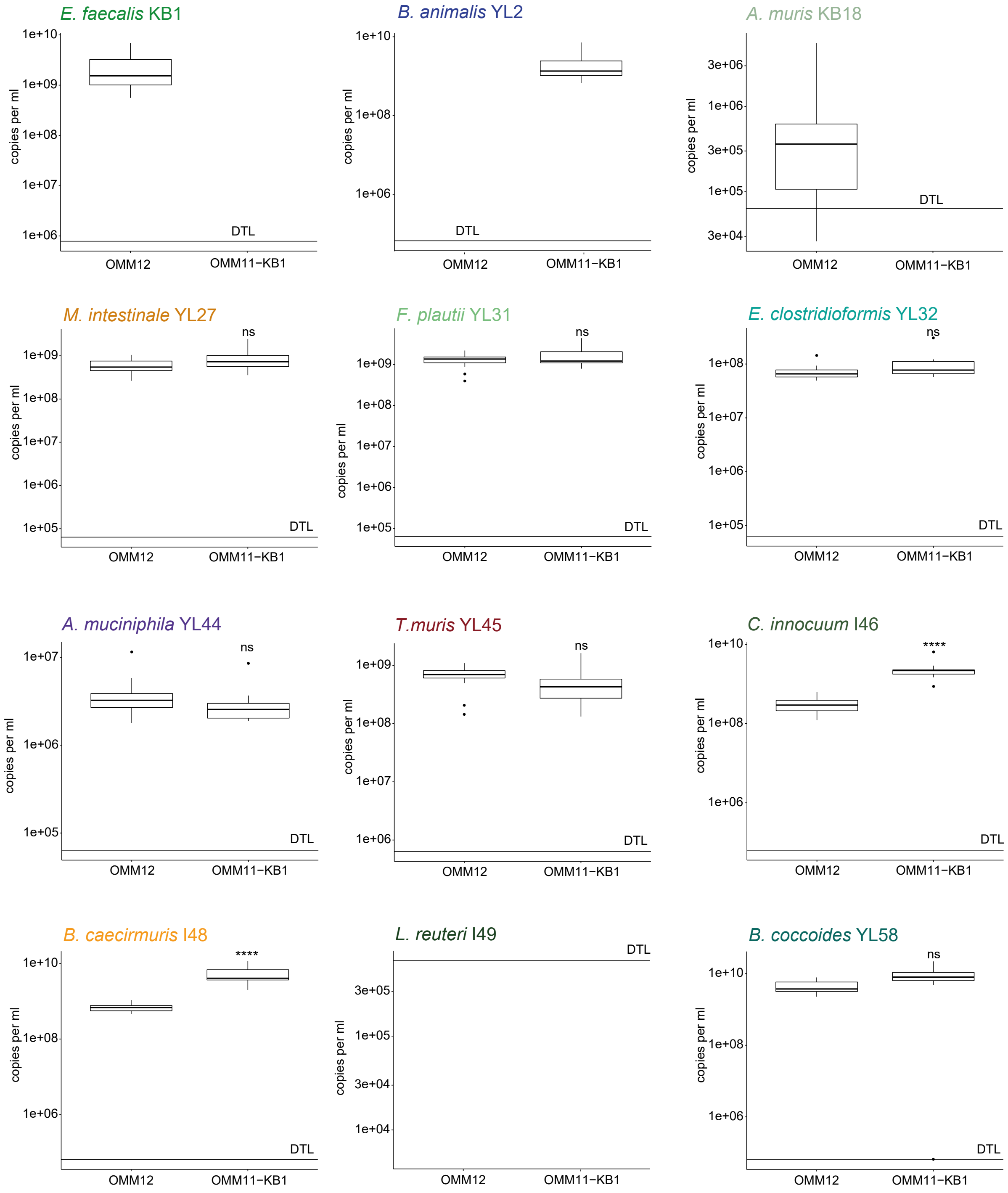


**Fig. S20. Comparison of absolute strain abundances in a full OMM$^{11\text{-KB1}}$ consortium vs. a dropout community.** Absolute abundance of strains was determined by qPCR as 16S rRNA copies per ml culture. The qPCR detection limit is marked with a horizontal line (DTL). Strain *L. reuteri* I49 was below detection limit in both communities in all replicates. Using a t-test absolute abundances were compared between the full community cultures (OMM$^{12}$, N=20) and a dropout community lacking *E. faecalis* KB1 (OMM$^{11\text{-KB1}}$, N=10). Significance levels are indicated as not significant (ns), p $\leq 0.05(*), p \leq 0.01(**), p \leq 0.001(***)$.

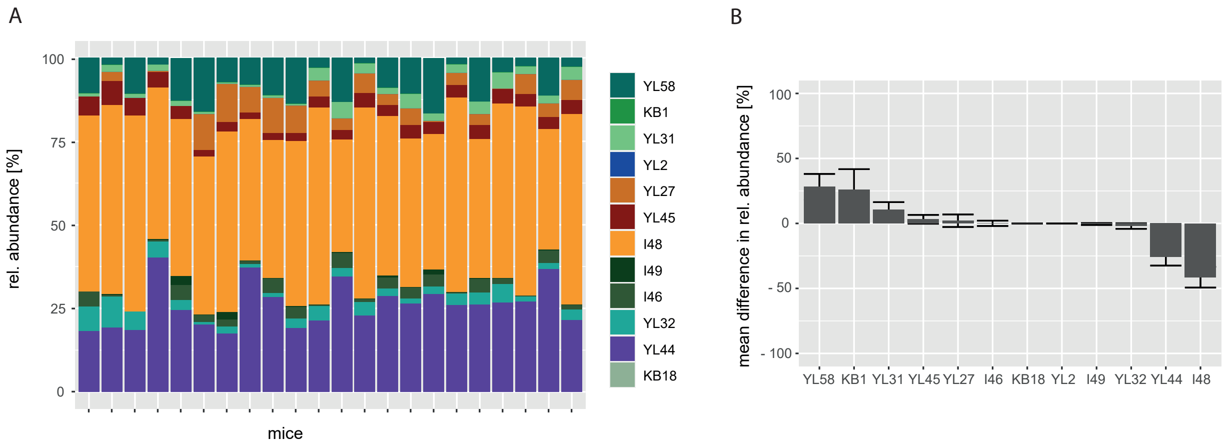


**Fig. S21. Relative abundance of the OMM$^{12}$ strains *in vivo*.** (**A**) Relative abundance of all twelve strains as determined by qPCR is shown for individual mouse feces samples (N=22). Data was taken from Eberl *et al.*[1]. (**B**) Analysing the mean difference between relative abundances *in vivo* (N=22) compared to *in vitro* batch culture (N=20) reveals that seven of twelve strains show a comparable relative abundance in both experimental settings. The corresponding standard deviation is shown as error bar.

[1] Eberl, C., Ring, D., Münch, P. C., Beutler, M., Basic, M., Slack, E. C., Schwarzer, M., Srutkova, D., Lange, A., Frick, J. S., Bleich, A., Stecher, B. (2020). Reproducible Colonization of Germ-Free Mice With the Oligo-Mouse-Microbiota in Different Animal Facilities. Frontiers in microbiology, 10, 2999. https://doi.org/10.3389/fmicb.2019.02999

#### Supplemental Tables

| **carbon source** | **area [%]** |
|---|---|
| Glucose | 54.42 |
| Fructose | 31.09 |
| Mannose | 3.62 |
| Galactose | 1.8 |
| Trehalose | 3.8 |

**Table S1. Analysis of carbon sources in AF-medium determined by GC-MS** Using internal standards, the listed sugars were detected in AF-medium. Relative concentrations of the individual sugars are depicted in percent of the total peak area as determined by GC-MS peak intensity.

| RT [min] | metabolite |
|---|---|
| 5 763 | Formate |
| 6 789 | Acetate |
| 8 336 | Propionate |
| 9 833 | Butyrate |
| 12 177 | Pyruvate |
| 15 509 | Lactate |
| 15 702 | Glycolate |
| 16 113 | Alanine |
| 16 372 | Glycine |
| 16 606 | $\beta$-Lactate |
| 17 443 | Urea |
| 17 552 | Valine |
| 17 672 | Norvaline |
| 18 014 | Leucine |
| 18 380 | Isoleucine |
| 18 662 | Succinate |
| 18 763 | Uracil |
| 18 818 | Proline |
| 20 459 | Phosphate |
| 20 712 | pyro-Glutamate |
| 20 892 | Methionine |
| 21 108 | Serine |
| 21 437 | Threonine |
| 22 103 | Phenylalanine |
| 22 655 | Aspartate |
| 23 141 | Cysteine |
| 23 331 | Hypoxanthine |
| 23 690 | Glutamate |
| 23 757 | Palmitate |
| 23 983 | Asparagine |
| 24 615 | Lysine |
| 25 429 | Stearat |
| 26 341 | Histidine |
| 26 516 | Citrate |
| 26 704 | Tyrosine |
| 27 079 | Tryptophan |

**Table S2. Analysis of metabolites in AF-medium determined by GC-MS** Using an internal norvaline standard, the listed metabolites were detected in AF-medium. The list of compounds is non-comprehensive.

| | mean GR [$h^{-1}$] | sd GR |
|---|---|---|
| *E. faecalis* KB1 | 2.40 | 0.11 |
| *B. animalis* YL2 | 1.74 | 0.06 |
| *A. muris* KB18 | 0.94 | 0.03 |
| *M. intestinale* YL27 | 1.28 | 0.04 |
| *F. plautii* YL31 | 1.13 | 0.02 |
| *E. clostridioformis* YL32 | 1.33 | 0.04 |
| *A. muciniphila* YL44 | 0.36 | 0.01 |
| *T. muris* YL45 | 0.78 | 0.02 |
| *C. innocuum* I46 | 1.83 | 0.08 |
| *B. caecimuris* I48 | 1.26 | 0.05 |
| *L. reuteri* I49 | 1.28 | 0.02 |
| *B. coccoides* YL58 | 1.64 | 0.06 |

**Table S3. Growth rates of the OMM[12] strains in monoculture.** Strain specific monoculture growth rates in AF-medium were determined by time-resolved measurements of OD at 600nm and linear fitting of the exponential growth phase. Strains were grouped by growth rate (GR) into fast growing strains (shown in blue, GR > 1.5 h-1), strains with intermediate growth rate (shown in black, GR > 1 h-1) and slow growing strains (shown in red, GR < 1 h-1).

### Supplemental Text A

##### Individual pH profiles as indicators for niche modification

The chemical composition of the individual SM is altered by substrate depletion and release of metabolic by- or waste products and change in pH. This change may lead to a different metabolic behavior of the strains while growing in SM. To quantify this, we determined the factor $\Delta\Delta$pH, which is the Euclidean distance between the $\Delta pH_{SM}$ that a strain shows after growth in fresh AF medium and the corresponding $\Delta pH_{DSM}$ after growth in each respective SM (**Fig. 3B and C**, **Fig. S5**). This factor, $\Delta\Delta$pH, can reflect an altered metabolic behavior of a strain in a specific chemical environment in another strain's SM. For example, after growth of *B. coccoides* YL58 in fresh AF medium the strain specific $\Delta pH_{SM}$ was found as $\Delta pH_{SM}$ = -1.11 ($pH_{SM,\ YL58}$ = 5.89). After growth of *B. coccoides* YL58 in the SM of *L. reuteri* I49 ($pH_{SM,\ I49}$ = 6.90) the strain specific $\Delta pH_{DSM}$ was found as $\Delta pH_{DSM}$ = -0.58 ($pH_{DSM}$ = 6.32), resulting in $\Delta\Delta$pH = 0.52. To identify cases, where products of a focal strain may lead to altered metabolic profiles of another strain, we correlated the $\Delta\Delta$pH values with the corresponding growth inhibition factor $d_{AUC}$ (**Fig. S5**). We reasoned, that the metabolic effect (e.g. change in metabolic profile of a strain growing in a specific SM compared to fresh medium) is most pronounced if the overall growth is little affected (rel. change in AUC $d_{AUC} > -0.4$), but the $\Delta\Delta$pH is large (Euclidean distance of pH profiles $\Delta\Delta pH > 0.5$). This was the case for *E. faecalis* KB1, *B. animalis* YL2, *M. intestinale* YL27, *B. cecimuris* I48 and *B. coccoides* YL58 in several different SM (**Fig. S5**, upper right quadrant). While these five strains were found to strongly acidify fresh culture medium during growth, they only weakly acidify the neutral SM of several other strains, including the SM of *A. muciniphila* YL44, *C. innocuum* I46 and *L. reuteri* I49. This suggests that the three latter strains alter their chemical environment in a way that somehow rewires metabolism of the others. The underlying changes may include the production of specific metabolic end products, e.g. SCFAs or depletion of favored growth substrates that require utilization of other metabolic pathways.

#### Supplemental Text B

**Genes for the production of antibacterial compounds by *E. faecalis* KB1**
The genome of *E. faecalis* KB1 (accession number CP022712.1) was screened for genes for the production of enterococcal bacteriocins (enterocins) using p-blast.

Query ID: **Enterocin L50A**
CP022712.1_2166 # 2333904 # 2334038 # -1 # ID=1_2166; partial=00; start_type=ATG; rbs_motif=GGAGG; rbs_spacer=5-10bp; gc_cont=0.281
Sequence ID: Query_66224 Length: 45
Range 1: 1 to 44
Score: 85.9 bits(211), Expect: 4e-26, Method: Compositional matrix adjust.,
Identities: 43/44 (98%), Positives: 44/44 (100%), Gaps: 0/44 (0%)
Query: MGAIAKLVAKFGWPIVKKYYKQIMQFIGEGWAINKIIEWIKKHI
Consence: MGAIAKLVAKFGWPIVKKYYKQIMQFIGEGWAINKII+WIKKHI
Subject: MGAIAKLVAKFGWPIVKKYYKQIMQFIGEGWAINKIIDWIKKHI

Query ID: **Enterocin L50B**
CP022712.1_2165 # 2333753 # 2333884 # -1 # ID=1_2165; partial=00; start_type=ATG; rbs_motif=AGGA/GGAG/GAGG; rbs_spacer=11-12bp; gc_cont=0.364
Sequence ID: Query_19555 Length: 44
Range 1: 1 to 43
Score: 83.2 bits (204), Expect: 4e-25, Method: Compositional matrix adjust.,
Identities: 40/43 (93%), Positives: 41/43 (95%), Gaps: 0/43 (0%)
Query: MGAIAKLVFKFGWPLIKKFYKQIMQFIGQGWTIDQIEKWLKRH
Consence: MGAIAKLV KFGWP IKKFYKQ+MQFIGQGWTIDQIEKWLKRH
Subject: MGAIAKLVAKFGWPFIKKFYKQVMQFIGQGWTIDQIEKWLKRH

Query ID: **Enterocin O16**
CP022712.1_26 # 23017 # 23529 # 1 # ID=1_26; partial=00; start_type=ATG; rbs_motif=AGxAGG/AGGxGG; rbs_spacer=5-10bp; gc_cont=0.366
Sequence ID: Query_4126 Length: 171
Range 1: 103 to 170
Score: 134 bits (338), Expect: 1e-43, Method: Compositional matrix adjust.,
Identities: 68/68 (100%), Positives: 68/68 (100%), Gaps: 0/68 (0%)
Query: LGSCVANKIKDEFFAMISISAIVKAAQKKAWKELAVTVLRFAKANGLKTNAI-IVAGQLAWAVQCGLSL
Consence: LGSCVANKIKDEFFAMISISAIVKAAQKKAWKELAVTVLRFAKANGLKTNAI-IVAGQLALWAVQCGLS
Subject: LGSCVANKIKDEFFAMISISAIVKAAQKKAWKELAVTVLRFAKANGLKTNAI-IVAGQLALWAVQCGLS

Query ID: **Enterocin 96**
CP022712.1_1870 # 2055051 # 2055275 # -1 # ID=1_1870; partial=00;start_type=ATG; rbs_motif=AGGAG; rbs_spacer=5-10bp; gc_cont=0.329
Sequence ID: Query_17682 Length: 75
Range 1: 27 to 74
Score: 102 bits (253), Expect: 4e-32, Method: Compositional matrix adjust.,
Identities: 48/48 (100%), Positives: 48/48 (100%), Gaps: 0/48 (0%)
Query: MSKRDCNLMKACCAGQAVTYAIHSLLNRLGGDSSDPAGCNDIVRKYCK

Consence: MSKRDCNLMKACCAGQAVTYAIHSLLNRLGGDSSDPAGCNDIVRKYCK
Subject: MSKRDCNLMKACCAGQAVTYAIHSLLNRLGGDSSDPAGCNDIVRKYCK

### Supplemental Text C

**Broad-scale draft metabolic models of the individual OMM$^{12}$ community members**

To allow a quick overview of the potential metabolic pathways employed by the OMM$^{12}$ bacteria, simplified metabolic models for individual bacterial species were constructed as follows: Depleted or produced metabolites were identified from metabolomics data. C5 and C6 sugars were assumed to be present in the gastrointestinal system, with selected C6 sugars identified by GC-MS (**Table S1**). $CO_2$ and molecular hydrogen were assumed as potential substrates and/or products. Since they are present in high concentration in the gas mix used for cultivation, changes in concentrations are not detectable. Potential metabolic connections were inferred from the substrate and product range and the genomic potential (**Fig. 3A, Fig. S9**) for major metabolic pathways of carbon compounds and amino acids. Pathways, their substrates and endpoints were identified from the Metacyc database (https://metacyc.org, January 2021). Pathways described for eukaryotes only were discarded.

The focus of our models lies mainly on the energy metabolism of the bacteria, excluding for example the synthesis and use of vitamins and secondary metabolites, as it is outside of the scope of this analysis. Similarly, we did not further investigate the synthesis of cellular material, as we presumed that it did not differ significantly between species and quantitative data is lacking. Therefore, also the utilization of hypoxanthine and uracil (both precursors for nucleotides) by some species has not been included in the models.

Most of the bacteria harbor a variety of potential pathways for energy metabolism, allowing them to access a range of substrates and thus compete in the community. Notable exceptions to this are *L. reuteri* I49 and *T. muris* YL45 with rather specialized metabolic profiles. However, their fates in the community are strikingly different: While *L. reuteri* I49 is not detectable in the community, *T. muris* YL45 can stably establish itself, possibly thanks to its sugar-independent energy generation using e.g. the Wood-Ljungdahl pathway. Further, similar to its close relative *Parasuterella excrementihominis* [2] it seems to be generally asaccharolytic, consume formate and acetate and produce succinate and lactate as fermentation end products. As *Parasutterella*, *T. muris* YL45 breaks down non-essential amino acids such as L-asparagine and L-aspartic acid. Furthermore, *T. muris* YL45 harbors the genes for dissimilatory nitrate reduction to ammonia (DNRA). Ammonia production may explain why growth of *T. muris* YL45 leads to an increase in pH.

However, since also species with a more diverse metabolism such as *A. muris* KB18 and *B. animalis* YL2 are not highly abundant, it is clear that the reasons for persistence in the community go beyond metabolic adaptability, encompassing for example bacteriocin production or efficient strategies for nutrient intake under competitive conditions.

[2] Ju, T., Kong, J. Y., Stothard, P., Willing, B. P. (2019). Defining the role of Parasutterella, a previously uncharacterized member of the core gut microbiota. The ISME journal, 13(6), 1520–1534. https://doi.org/10.1038/s41396-019-0364-5
